# Targeting the trypanosome kinetochore with CLK1 protein kinase inhibitors

**DOI:** 10.1101/616417

**Authors:** Manuel Saldivia, Srinivasa P.S. Rao, Eric Fang, Elmarie Myburgh, Elaine Brown, Adam J. M. Wollman, Ryan Ritchie, Suresh B Lakhsminarayana, Yen Liang Chen, Debjani Patra, Hazel X. Y. Koh, Sarah Williams, Frantisek Supek, Daniel Paape, Christopher Bower-Lepts, Mark C. Leake, Richard McCulloch, Marcel Kaiser, Michael P. Barrett, Jan Jiricek, Thierry T. Diagana, Jeremy C. Mottram

## Abstract

The kinetochore is a macromolecular structure that assembles on the centromeres of chromosomes and provides the major attachment point for spindle microtubules during mitosis. In *Trypanosoma brucei* the proteins that make up the kinetochore are highly divergent, with the inner kinetochore comprising at least 20 distinct and essential proteins (KKT1-20) that include four protein kinases, CLK1 (KKT10), CLK2 (KKT19), KKT2 and KKT3. We performed a phenotypic screen of *T. brucei* bloodstream forms with a Novartis kinase-focused inhibitor library, which identified a number of selective inhibitors with potent pan-kinetoplastid activity. Deconvolution of an amidobenzimidazole series using a selection of 37 *T. brucei* mutants that over-express known essential protein kinases identified CLK1 as the primary target. Biochemical studies show that the irreversible competitive inhibition of CLK1 is dependent on a Michael acceptor forming an irreversible bond with C215 in the ATP binding pocket, a residue that is not present in human CLK1, thereby providing selectivity. Chemical inhibition of CLK1 impairs inner kinetochore recruitment and compromises cell cycle progression, leading to cell death. We show that KKT2 is a substrate for CLK1 and identify phosphorylation of S508 to be essential for KKT2 function and for kinetochore assembly. We propose that CLK1 is part of a novel signalling cascade that controls kinetochore function via phosphorylation of the inner kinetochore protein kinase KKT2. This work highlights a novel drug target for trypanosomatid parasitic protozoa and a new chemical tool for investigating the function of their divergent kinetochores.

---

Nearly a billion people in over 80 countries are at risk for one of the three kinetoplastid diseases: leishmaniasis, Chagas disease, and human African trypanosomiasis (HAT). While there have been notable advances, more than 30,000 people continue to die each year from complications of these neglected diseases and millions more suffer from serious morbidities. Current therapies have severe shortcomings. There is an urgent need for novel treatments that are safe, efficacious and easy to administer. As part of a phenotypic screen of ∼2.2 million compounds against *T. brucei* mammal-infective bloodstream forms^1^ we used the Novartis kinase focused inhibitor library to identify additional growth inhibitors of the parasite. This led to the identification of AB0, an amidobenzimidazole (Supplementary figure 1a and Fig. 1a) with an EC_50_ of 230 nM in an *in vitro T. brucei* cell growth assay (Fig. 1a). Detailed structure activity relationship (SAR) studies with over 230 compounds shows that the introduction of a Michael acceptor chemical functionality is critical for potent activity and affords higher lipophilic efficiency (LipE >3) required for more favourable drug-like properties (Fig. 1b). SAR analysis also led to the identification of more potent bioactive compounds, including AB1 (EC_50_ of 72 nM against *T. brucei*), with a >100-fold cytotoxicity window (Fig. 1). AB1 not only showed potent activity against *T. b. gambiense* and *T. b. rhodesiense*, the causative agents of sleeping sickness but also showed pan-kinetoplastid activity against *T. cruzi* (aetiology for chagas disease), *Leishmania mexicana* (cutaneous leishmaniasis) *and L. donovani* (visceral leishmaniasis) suggesting that the molecular target is conserved across trypanosomatids (Fig. 1a). AB1 compound was cidal to *T. brucei* showing concentration and time dependent kill, further it also exhibited relapse free cidality in wash-off assays suggesting ability to achieve sterile cure (Supplementary Fig 1b,c). AB1 had better solubility and *in vitro* pharmacokinetic properties with moderate to low clearance in mice, rat and human microsomes, allowing us to evaluate the ability to cure HAT in mouse models (Fig. 1a). *In vivo* efficacy of AB1 was tested in mice with haemolymphatic and central nervous system (CNS) *T. brucei* infections using an optimised bioluminescence HAT mouse model ^2^. A dose dependent cure was observed with AB1, resulting in relapse free cure with a daily oral dosage of 50 mg/kg for four consecutive days in a haemolymphatic mouse model of infection (Fig. 1c). However, despite a 4 log reduction in parasitaemia observed in a pleomorphic GVR35 strain chronic (CNS) model of infection, the parasites relapsed, most likely due to poor access of AB1 to parasites in the brain (Supplementary Fig. 2a,b). Although AB1 showed good brain partitioning due to high brain tissue binding (>99%), the free fraction available for acting against the parasites in the brain was negligible, leading to poor efficacy (Supplementary Fig. 2c,d). Further medicinal chemistry optimization is required to find compounds with better pharmacokinetic properties in order to achieve cure in the CNS model. The mouse efficacy studies using AB1 showed *in vivo* chemical validation of AB series compounds as promising anti-trypanosomatid candidates.

**Fig. 1.**
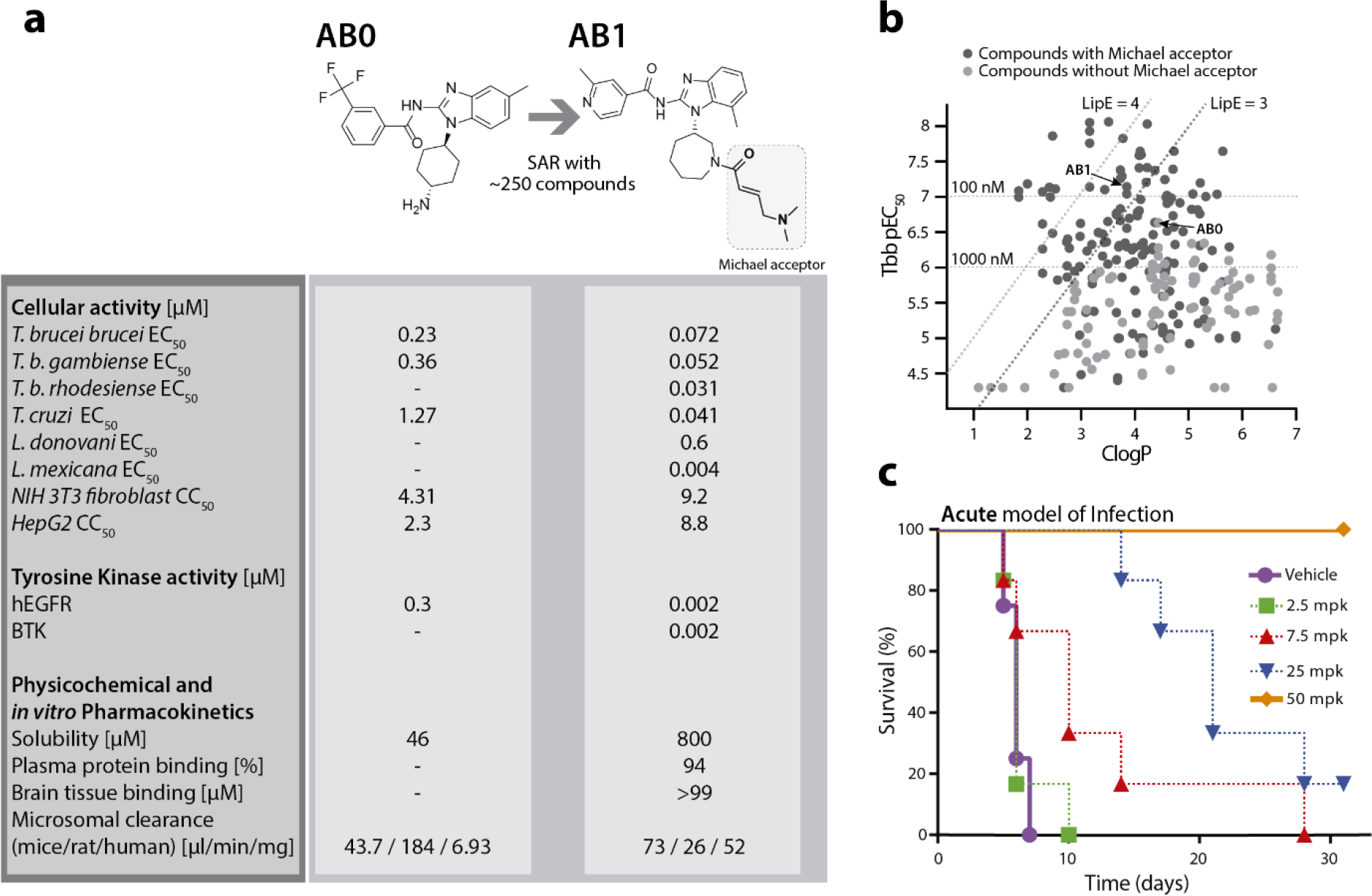
Pan kinetoplastid activity of amidobenzimidazole (AB) series. (a) Structure, anti-kinetoplastid parasite activity, cytotoxicity, tyrosine kinase inhibition and *in vitro* pharmacokinetics of AB0 and AB1.
(b) Structure activity relationship of amidobenzimidazoles. Compounds with the Michael acceptor were more potent against *T. brucei* (Tbb EC_50_) and had better lipophilic efficiency (clogP, calculated octanol-water partition coefficient) compared to non-Michael acceptor compounds.
(c) *In vivo* activity of AB1 in a haemolymphatic mouse model for human African trypanosomiasis. Note the dose dependent activity of AB1, achieving complete cure at 50 mg/kg once daily dose.

**Fig 2.**
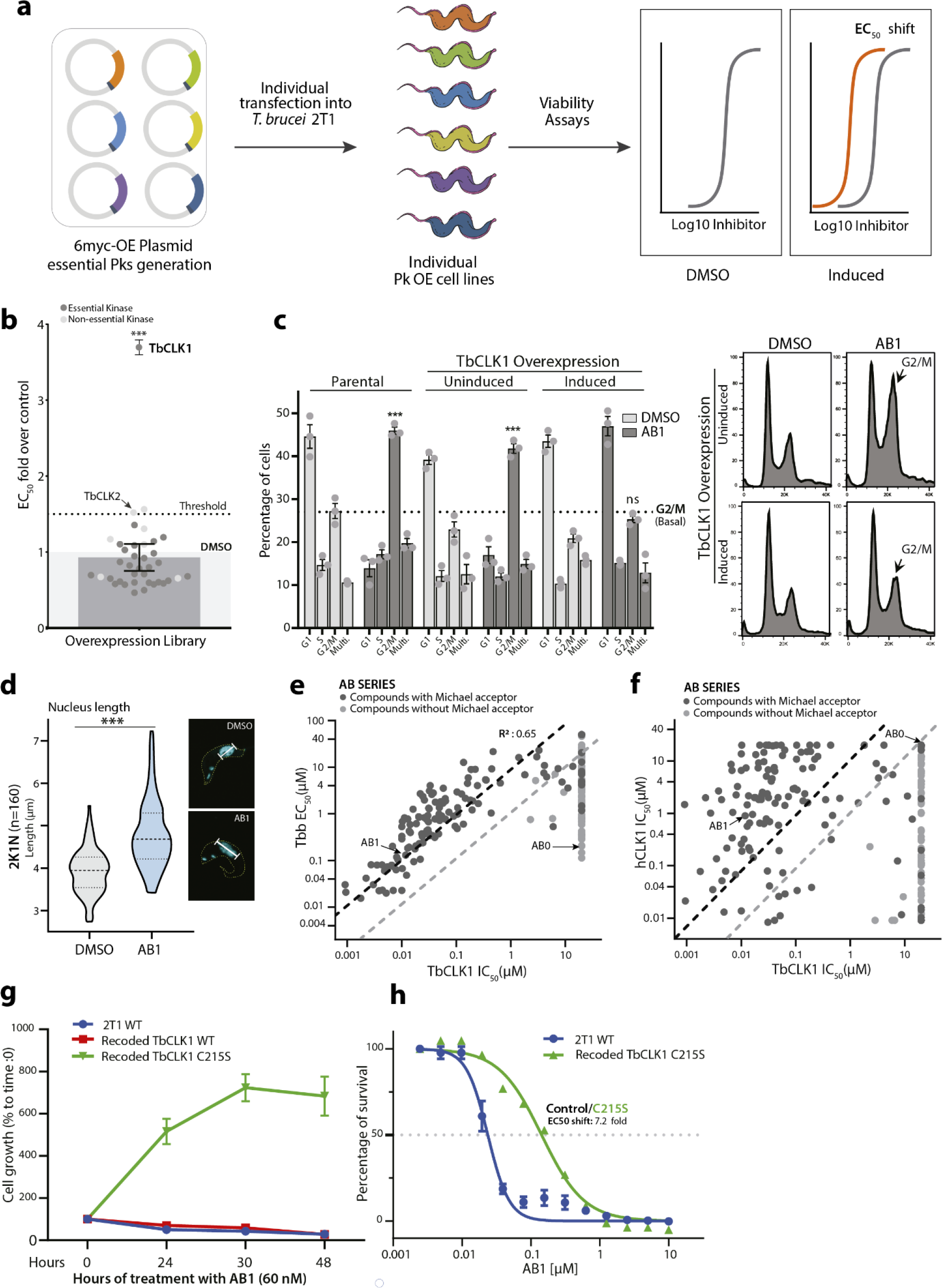
Identification of TbCLK1 as the molecular target for the AB series. (a) Inducible overexpression (OE) of essential protein kinases (Pks) and target deconvolution approach for the AB series. Schematic representation of the experimental workflow: 6-myc OE plasmids, each containing a specific protein kinase tagged with six myc epitopes, were individually transfected into 2T1 bloodstream form *T. brucei*. Viability of induced individual *T. brucei* OE lines after treatment was assessed by measuring the conversion of resazurin (Alamar blue) to resorufin.
(b) AB1 half maximal effective concentration (EC_50_) was analysed from 37 individual protein kinase OE cell lines, including essential (dark grey circles) and non-essential control (light grey circles) protein kinases. The graph represents EC_50_ fold change over the parental cell line (2T1 cell line, DMSO) (light grey shade box), where overexpression of CLK1 confers resistance to AB1. Error bars show the SEM. Dark grey bar represents the EC_50_ average of all cell lines. P values were calculated using a Two-tailed Student’s t-tests comparing TbCLK1 OE with parental cell line where *** p-value <0.001. TbCLK2 (a non-essential kinase) EC_50_ (dotted line in the graph) was chosen as the threshold, due its structural similarity with TbCLK1.
(c) TbCLK1 OE confers resistance to AB1-induced cell cycle arrest. TbCLK1 OE was induced or not with tetracycline for 18 hr, and cells then incubated for 6 hr with 5 x AB1 EC_50_ (dark grey bars); the 2T1 cell line was a parental control. Cell cycle distribution was determined by flow cytometry. Left: dotted line represents (basal) G2/M untreated average and *** represents a p-value <0.001 comparing untreated vs treated cell lines. Right: cell cycle profile histogram of cells stained with propidium iodide showing G2/M cell cycle accumulation (arrow). P values were calculated using a Two-tailed Student’s t-tests.
(d) Violin plot of average length (µm) of the nuclei from parasites treated (blue) or not (grey) with AB1 (n=160 2K1N parasites, *** p-value <0.001). Right: Example of a cell stained with DAPI (cyan) from each condition.
(e) Positive correlation between inhibition of recombinant TbCLK1 enzyme (IC_50_) and growth inhibition (EC_50_) of *T. brucei* bloodstream form parasites with AB series compound (n=230; R^2^= 0.65).
(f) Lack of correlation between inhibition of recombinant human CLK1 and recombinant TbCLK1 enzyme (IC_50_) with AB compound series (n=230 compounds). Majority of compounds showed > 10 fold selectivity against TbCLK1 compared to hCLK1.
(g) TbCLK1 C215S mutant impairs the effect of AB1 on parasite growth. Recoded TbCLK1 (red), TbCLK1 C215S (green) and parental 2T1 cell line (blue) (1 × 10^5^ parasites ml^-1^) were treated with 60 nM AB1, and the percentage of cell growth calculated daily and compared to Day 0. Error bars represent mean ± SEM of three replicates.
(h) Recoded re-expression of TbCLK1 C215S confers an EC_50_ shift of resistance compared to parental cell line. Endogenous TbCLK1 C215S-HA mutant was expressed in the TbCLK1 RNAi line and AB1 half maximal effective concentration (EC_50_) was determined after 48 hr. Error bars represent mean ± SEM of three replicates.

AB1 is a known inhibitor of mutant human epidermal growth factor receptor (EGFR), a well-studied target for non-small cell lung cancer (Fig 1a) ^3^. Whilst EGFR belongs to the tyrosine kinase family, the *T. brucei* genome lacks members of the receptor-linked or cytosolic tyrosine kinase families ^4^. A major challenge for phenotypic hits is to identify the molecular target that is responsible for that effect. Since AB series compounds were hypothesised to target protein kinases, we generated inducible *T. brucei* gain-of function mutants of individual protein kinases to screen for resistance to AB1 (Fig. 2a). Thirty-seven essential protein kinases that had been assessed previously by RNAi as having specific RNAi induced cell cycle defects ^5^ were analysed, with six non-essential protein kinases used as controls. Only one protein kinase, CLK1, showed a significant 3.5 fold increase in resistance compared with its uninduced control (Fig. 2b). Overexpression of CLK2, a non-essential protein kinase closely related to CLK1 (92% sequence identity, Supplementary Fig. 3a), did not alter the resistance profile to AB1. CLK1 is conserved in different kinetoplastid parasites (Supplementary Fig. 3b).

**Fig. 3.**
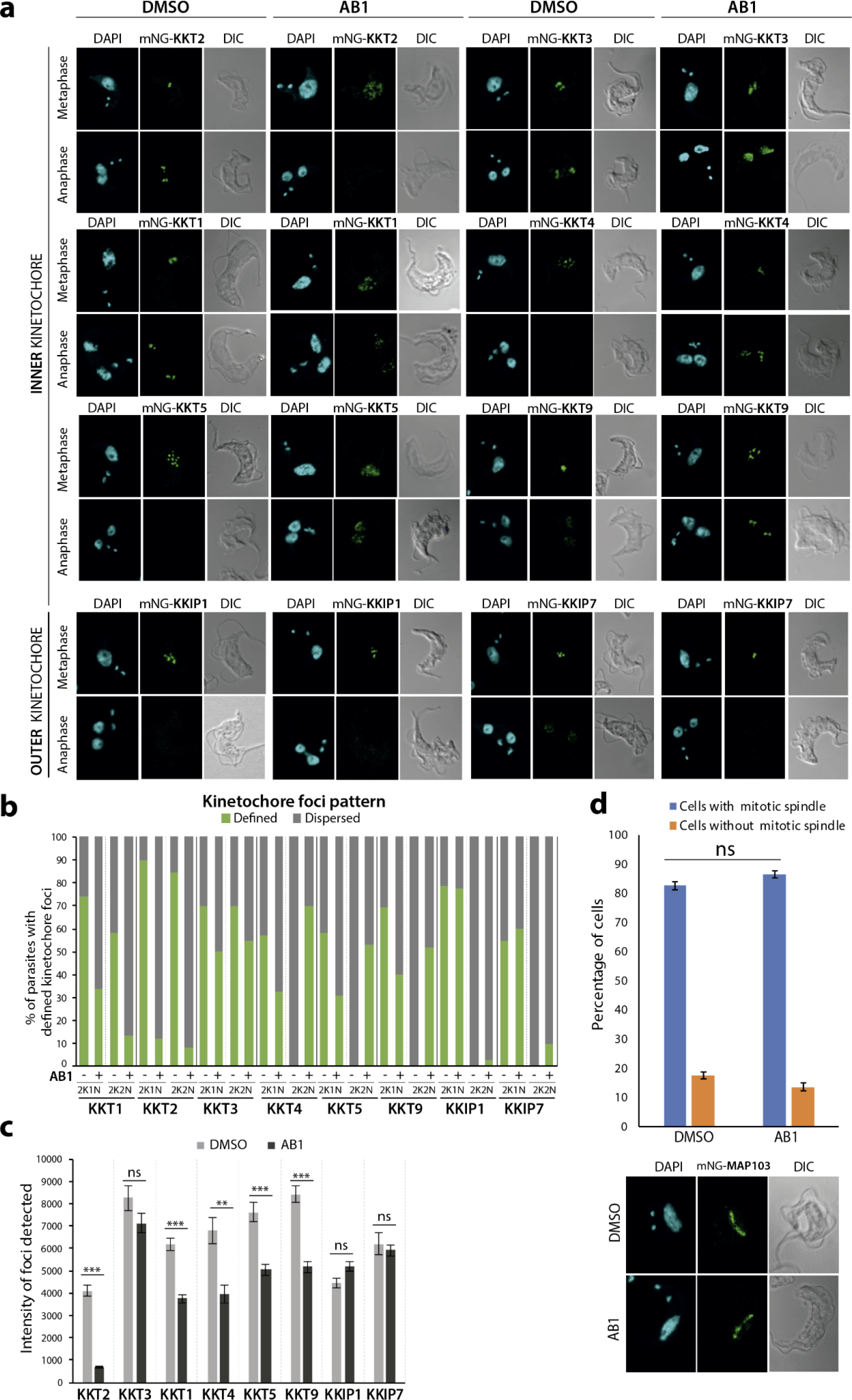
TbCLK1 inhibition impairs inner kinetochore dynamics. (a) Localization of inner (top panel) and outer (lower panel) kinetochore core components after TbCLK1 inhibition by AB1. Parasites were incubated or not for 6 hr with 5x EC_50_ AB1. Representative fluorescence microscopy micrographs, showing bloodstream form parasites endogenously expressing N-terminal mNeonGreen (mNG) tagged kinetochore proteins. Cells in metaphase and anaphase are shown. Cells were counterstained with DAPI to visualize DNA (cyan). The right panel shows the Nomarsky (DIC) corresponding images.
(b) Percentage of cells in metaphase (2K1N) and anaphase (2K2N) showing a defined kinetochore localisation before and after AB1 treatment as in (a) (n>100 cells in each stage).
(c) Intensity of KKT foci detected before (DMSO) and after AB1 treatment. The data represents 75% percentile of total foci intensity (n=80 kinetochores in each condition). Error bars, SEM; ** p<0.01, ***p < 0.001. ns not significant. (Mann–Whitney U test).
(d) Spindle formation after TbCLK1 inhibition. Parasites expressing mNG-MAP103 were left untreated or treated for 6 hr with 5x AB1 and analysed by confocal microscopy. Error bars, SEM (n>100 cells in metaphase). ns not significant. <u>Bottom:</u> representative micrograph of each condition.

In mammals, CLK1 belongs to the Clk (Cdc2-like kinase) family implicated in RNA splicing control and consists of at least four members ^6^. In *T. brucei*, CLK1 is a kinetochore component essential for mitosis and has been proposed to be one of the potential targets for the fungal antibiotic hypothemycin ^7–9^. As treatment with AB1 resulted in a G2/M cell cycle arrest (Fig. 2c) with most of the treated cells having an enlarged nucleus (Fig. 2d) similar to TbCLK1 RNAi knockdown, (Supplementary Figs. 4a-c), we tested if overexpression of TbCLK1 would confer resistance to drug-induced G2/M cell cycle arrest. Indeed, parasites overexpressing TbCLK1 had a normal cell cycle profile after treatment with AB1, in comparison to the parental cell line and DMSO uninduced control, which were G2/M arrested after treatment (Fig. 2c). TbCLK1 overexpression impairs parasite fitness by affecting cell cycle progression, suggesting that *T. brucei* tightly regulates CLK1 expression (Supplementary Fig. 4d). In addition, AB1 attenuates TbCLK1 toxicity, providing further evidence that CLK1 is the compound’s target (Supplementary Figs. 4e, f).

**Fig. 4.**
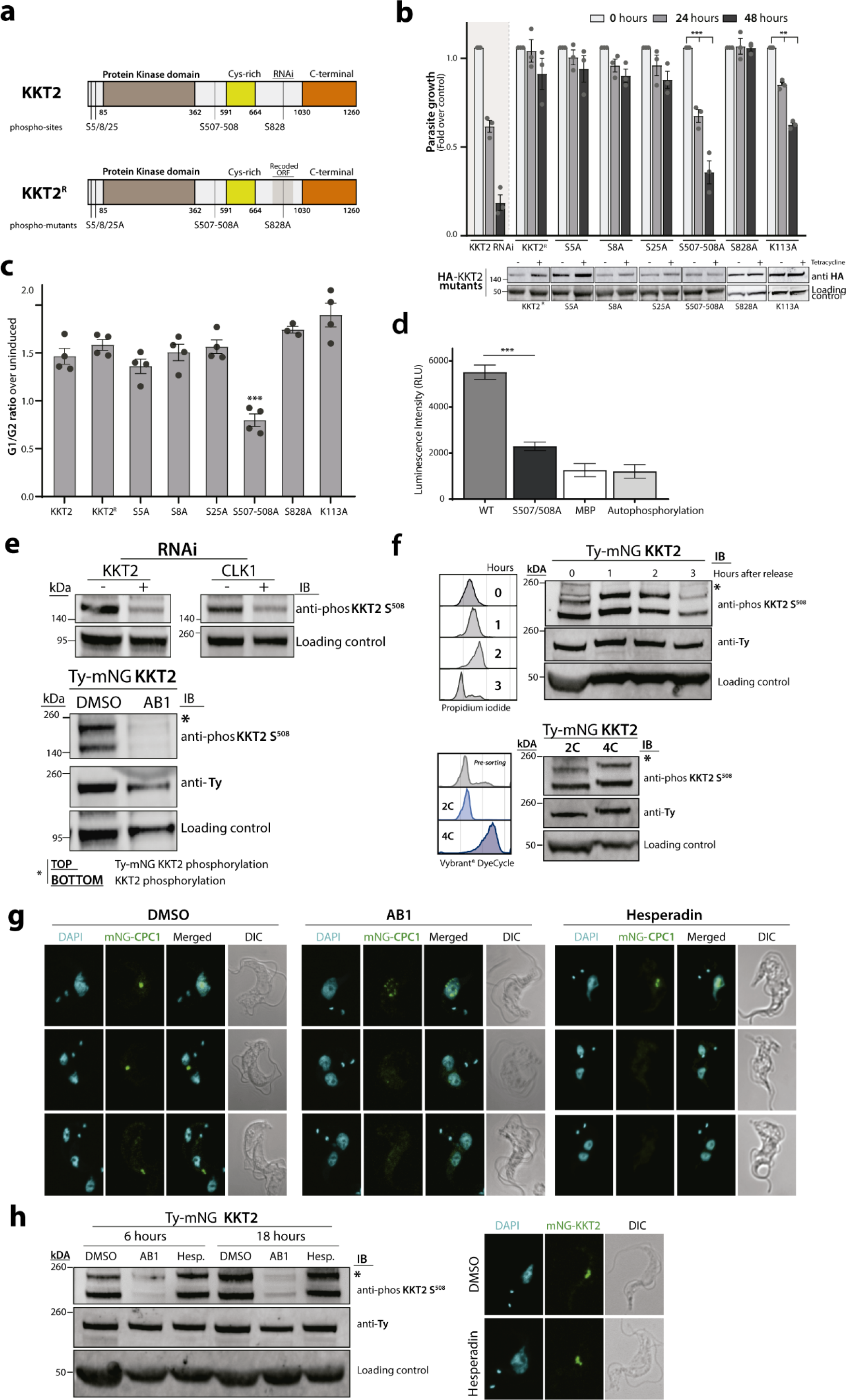
TbCLK1 regulates KKT2 function by phosphorylation of residue S508. (a) Schematic representation showing known KKT2 phosphosites.
(b) *In vitro* growth profile of KKT2 RNAi, KKT2^R^ and KKT2^R^ phosphomutants and active site mutant. Bars showing cumulative fold over uninduced control counts over time following tetracycline induction of cell lines in culture. Error bars, SEM; ** p<0.01, ***p < 0.001. <u>Lower</u> <u>panel:</u> Western blot of HA-KKT2 mutants. The expression of KKT2 phosphomutants mutants were detected using an anti-HA antibody. EF-1 alpha protein expression was used as the loading control.
(c) Cell cycle profile of KKT2 RNAi, KKT2^R^ and KKT2^R^ phosphomutants. Bars showing G1/G2 ratio over uninduced control following tetracycline induction of cell lines in culture. Error bars represeent SEM of 4 replicates. P values were calculated using a Two-tailed Student’s t-tests; where ***p < 0.001.
(d) Recombinant CLK1 phosphorylates recombinant KKT2 *in vitro*. Recombinant fragment of KKT2 including S^507-S508^ (KKT2^486-536^) was used as substrate for rTbCLK1 by ADP-Glo™ Kinase Assay. Same fragment, but including a S^507A-S508A^ mutation was used as a control. Phosphorylation of maltose binding protein (MBP) and rTbCLK1 autophosphorylation (not substrate) was included as control. Error bars, SEM; ***p < 0.001 (Two-tailed Student’s t-test).
(e) Specificity of KKT2 S^508^ phospho-specific antibody. <u>Top:</u> CLK1 and KKT2 RNAi was induced in 2T1 parasites for 24 hr. KKT2 phosphorylation was analysed by western blot using KKT2 S^508^ phospho-specific antibody. Expression of an endogenous protein and Ty-KKT2 were used as KKT2 and CLK1 RNAi loading control respectively. <u>Bottom:</u> Phosphorylation of KKT2 S^508^ and Ty-mNG KKT2 S^508^ after 18 hr 2× EC_50_ AB1 treatment. Expression of Ty-KKT2 and EF-1 alpha protein expression were used as the loading control.
(f) KKT2 S^508^ phosphorylation during the cell cycle. <u>Top:</u> cells expressing Ty-mNG-tagged KKT2 were synchronized in late S phase by incubating with 10 µM hydroxyurea for 6 hr and released. After release, cells were collected after 0, 1, 2 or 3 hr and KKT2 S^508^ phosphorylation was analysed by western blot. Cell cycle progression was assessed by flow cytometry (left) by staining with propidium iodide. <u>Bottom</u>: KKT2 S^508^ phosphorylation profile of post-sort 2C and 4C populations by flow cytometry Vybrant DyeCycle Violet stained Ty-mNG-tagged KKT2 cell line. Post-sort efficiency of each population is shown in the left panel.
(e) Localisation of TbCPC1 after treatment with AB1 or Hesperadin. Ty-mNG-TbCPC1 expressing parasites were left untreated or treated for 6 hr with 250 nM AB1 or 800 nM Hesperadin and analysed by confocal microscopy. Cells in metaphase and anaphase are shown. Cells were counterstained with DAPI to visualize DNA (cyan). The right panel shows the Nomarsky (DIC) corresponding images.

We expressed recombinant TbCLK1 and human CLK1 and tested over 230 AB compounds in the series to determine the biochemical structure-activity relationship. A strong correlation was observed between inhibition of the *T. brucei* CLK1 enzyme and cellular activity (R^2^ = 0.65), which supports the chemical validation of CLK1 as the molecular target for amidobenzimidazoles (Fig. 2e). The majority of the compounds showed greater selectivity against TbCLK1 compared to hCLK1 (Fig. 2f). AB1 inhibited TbCLK1 with an IC_50_ of 10 nM, with 90-fold selectivity observed over hCLK1 (Fig. 2f). Further, only compounds with a Michael acceptor inhibited CLK1 kinase activity, strongly suggesting this functionality is an essential feature of the pharmacophore (Fig. 2e). In order to test the putative critical role played by the Michael acceptor, a compound similar to AB1, which has a saturated double bond at the Michael acceptor, was profiled in both enzymatic and cellular assays. This compound (AB2) was completely inactive in both TbCLK1 enzyme (IC_50_ > 20 µM) and whole cell growth inhibition assays (Tbb EC_50_ > 50 µm), confirming the importance of the Michael acceptor for activity. The human EGFR inhibitors having Michael acceptors are known to covalently interact with an active site cysteine (C797) near the ATP binding domain ^3, 10^. In order to determine the probable TbCLK1 cysteine interacting with AB1, the binding mode of AB1 was modelled in silico using the crystal structure coordinates of hCLK1 (pdb code: 1z57) overlaid with the co-crystal of hEGFR:AB1 (pdb code: 5feq). Subsequent sequence alignment between hCLK1 and TbCLK1 identified C215 to be the probable covalent cysteine and an explanation for the selectivity, since this residue is a serine in hCLK1 (Supplementary Fig. 5a). To investigate the covalent binding and interaction with C215, we co-incubated AB1 and AB2 with both wild type and C215A mutant of TbCLK1 and assessed by mass-spectrometry. As expected, incubation of AB1 with wild type TbCLK1 enzyme resulted in one main product per protein which has average mass 475 Da higher than the unmodified protein (Supplementary Fig 5b). This was consistent with addition of one molecule of AB1, which has a mass of 475 Da. No shift in the mass was seen for AB2 an analog of AB1 with saturated Michael acceptor and in C215A mutant CLK1 (Supplementary Fig 5b, c). These results clearly indicated covalent interaction of Michael acceptor with C215 of TbCLK1. Further, to assess the importance of *T.brucei* CLK1 C215 in AB1 binding in parasites, we expressed a recoded C215A and C215S mutants in the TbCLK1 RNAi line and examined its effects on AB1 binding or parasite fitness. As shown in Supplementary figure 5d, TbCLK1 C215S expressing parasites had a normal cell cycle profile whilst C215A mutation triggers a G2/M cell cycle arrest, suggesting this residue is important for CLK1 function and therefore mimicking AB1 –TbCLK1 mediated inhibition. Thus, we evaluated if C215S mutation impairs AB1 binding, and as shown in Fig. 2g,h, this mutation conferred a 7-fold EC50 shift of resistance against AB1 and protected parasites from 48 hours of treatment with a sub-lethal concentration of AB1 (2 × EC50).

**Fig. 5.**
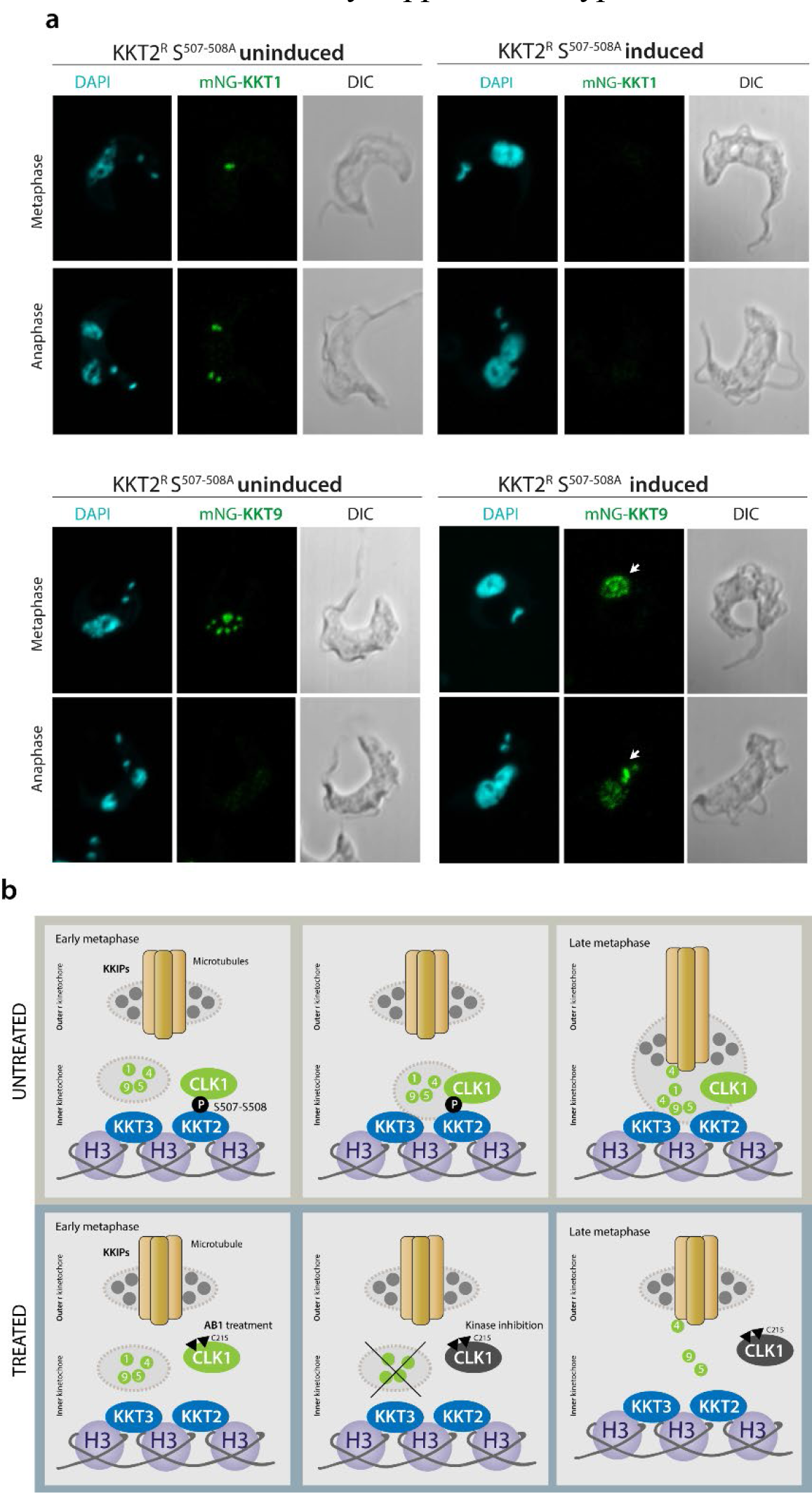
Phosphorylation of KKT2 is required for kinetochore assembly. (a) Recruitment of KKT1 and KKT9 to the kinetochore is impaired in KKT2^R^ S^507-508A^ mutant. Representative fluorescence microscopy of BSF parasites endogenously expressing KKT1 or KKT9 tagged with mNeonGreen (mNG) at the N terminus in metaphase and anaphase. Cells were imaged 48 hr after induction of the KKT2^R^ S^507-508A^ mutant. Cells were counter stained with DAPI to visualize DNA (cyan).
(b) Model for TbCLK1 function in kinetochore assembly. This schematic diagram summarizes the recruitment defects caused by inhibition of CLK1 by AB1. In untreated cells in early metaphase (top panel), CLK1 phosphorylates KKT2, resulting in recruitment of inner kinetochore components, allowing posterior kinetochore assembly to outer kinetochore components. Once CLK1 is inhibited by AB1 (lower panel), phosphorylation of KKT2 S^508^ is prevented, leading to a failure of recruitment of inner kinetochore components and consequent cell cycle arrest. H3, histone H3

TbCLK1 (KKT10) is a core component of the inner kinetochore. In other organisms, the kinetochore supports directional movement of chromosomes into microtubules to ensure faithful chromosome segregation ^11^. Some of the *T. brucei* kinetochore components have been recently described, and grouped according to their patterns of expression/localization through the cell cycle ^9, 12^. However, the underlying mechanisms behind the assembly of the kinetochore remain unclear. The significant divergence between components of the human and parasite kinetochores underpins its potential druggability. Although depletion of TbCLK1 has been associated with the presence of lagging chromosomes during mitosis ^9^ the causal link of TbCLK1 to this process has yet to be established. To investigate this further *T. brucei* bloodstream form cells were synchronised by cell sorting ^13^ to give 2C and 4C nuclear DNA content populations and then allowed to enter the cell cycle in the presence or absence of AB1 (Supplementary Fig. 6). Inhibition of TbCLK1 in the 2C population (G1-phase) synchronously progressing through the cell cycle led to arrest of cells in late metaphase, while inhibition of TbCLK1 in the 4C population synchronously progressing through the cell cycle induced an arrest in late anaphase. This suggests a dual role for TbCLK1 in promoting kinetochore assembly and movement of the chromosomes on the spindle during metaphase and exit from anaphase into cytokinesis.

Given the importance of kinetochore movement during metaphase in eukaryotes ^14^, we next assessed the impact of TbCLK1 activity on kinetochore dynamics using AB1 as a chemical tool in bloodstream form parasites. The expression and localisation of kinetochore proteins, tagged with mNeonGreen, were assessed by confocal microscopy. Consistent with previous observations in procyclic form cells ^9, 12^, we observed in bloodstream form cells different kinetochore timings and patterns of expression throughout the cell cycle; KKT2 and KKT3 are constitutively expressed until anaphase, whilst KKT1 gradually loads from S phase onwards until the end of mitosis, whilst KKT4, KKT5, and KKIP1 expression is restricted to metaphase. Furthermore, KKT9 and KKIP7 expression diminishes during anaphase, suggesting both proteins may be acting as scaffolds for the recruitment of multiple other components. Treatment with 5 x EC_50_ of AB1 caused dispersal, to varying degrees, for KKT1, KKT2, KKT5, KKT9, KKT13, KKT14 and KKT20 from the defined foci of the kinetochore within the nucleus, whilst KKT3, KKT7, KKT11, KKIP1 and KKIP7 remained in distinct foci (Fig. 3a,b and Supplementary Fig 7a). Automated foci detection using sub-pixel precise single particle localization combined with image segmentation ^15^ and intensity quantification ^16^ determined that there was a significant reduction in foci intensity for KKT1, KKT2, KKT4, KKT5 and KKT9, but not KKT3 (Fig. 3c and Supplementary Figs 7b, c). These results suggest that although KKT2 and KKT3 are centromere-anchored proteins, they might have different functions during kinetochore assembly, and that TbCLK1 is a key regulator of inner kinetochore component dynamics. Faithful chromosome segregation relies on the interaction between chromosomes and dynamic spindle microtubules ^17^. Our results suggest that localisation/expression of outer kinetochore proteins KKIP1 and KKIP7 remains unaffected after AB1 treatment, whereas KKT4, recently described as a microtubule tip–coupling protein ^18^, remains in anaphase, suggesting end-on interaction defects of microtubules with kinetochores. Spindle elongation is important for correct segregation of chromosomes during anaphase ^19^. To further examine if TbCLK1 inhibition impairs microtubule spindle dynamics, we analysed the expression of microtubule-associated protein 103 kDa (MAP103) ^20^ and showed that treatment with AB1 does not affect microtubule spindle formation (Fig. 3d). Considering that TbCLK1 inhibition during metaphase results in an arrest in late anaphase, it is likely that the function of TbCLK1 during cytokinesis is related to either the control of kinetochore-spindle microtubule attachment errors, or its interactions with the chromosomal passenger complex (CPC). Of note, it has been reported that *T. brucei* aurora kinase B has an important role during metaphase-anaphase transition and the initiation of cytokinesis via regulation of the CPC ^21–23^ and nucleolar and other spindle-associated proteins (NuSAPs) ^24^.

KKT2 and KKT3 protein kinases are likely components of the trypanosome inner kinetochore with functional equivalence to the constitutive centromere-associated network (CCAN), a canonical component of the eukaryotic inner kinetochore ^25^. It has been reported that phosphorylation of kinetochore proteins has critical roles in kinetochore organization and interaction during mitosis ^26^. We speculated that KKT2 provides a platform on which the kinetochore multi-protein complex assembles, and that phosphorylation orchestrates this process. To address whether KKT2 might be a TbCLK1 substrate, we first analysed mobility shifts of phosphorylated forms of KKT2 and KKT3 using Phos-tag™ gel electrophoresis ^27^. A low-mobility, non-phosphorylated form of KKT2 was detected after treatment with AB1 or after TbCLK1 depletion by RNAi, whilst KKT3 remained unaffected (Supplementary Fig. 8). Six phosphorylation sites have been identified in KKT2 (S^5^, S^8^, S^25^, S^507^, S^508^, S^828^) ^28^ and we tested if these are important for KKT2 function by generating a KKT2 RNAi line with a recoded HA-tagged version of KKT2 integrated into the ribosomal locus (Fig. 4a). This constitutively expressed KKT2 (*KKT2^R^*) is not susceptible to RNAi-mediated degradation and *KKT2^R^* complements the loss of function of KKT2 48 hrs after RNAi induction (Fig. 4b). Replacement of Ser for Ala in KKT2 at positions S^5^, S^8^, S^25^ and S^828^ resulted in complementation of KKT2 function when expressed in the RNAi line. In contrast, dual replacement of the KKT2 phosphorylation sites S^507^ and S^508^ with Ala (KKT2^S507A-S508A^) failed to complement loss of KKT2 function with respect to parasite growth (Fig. 4b) or cell cycle progression after 48 hours induction (Fig. 4c), despite good expression levels in the cell (Fig 4b. lower panel). These defects phenocopy the effect of AB1 and show the importance of the two phosphorylation sites for the function of KKT2. To assess whether protein kinase activity is essential for KKT2 function an active site mutant was generated in KKT2^R^ (KKT2^K113A^). A significant loss of function was observed after 48 hrs induction, indicating protein kinase activity is essential for KKT2 function, but not for regulating cell cycle progression (Fig. 4c).

To address whether TbCLK1 phosphorylates KKT2 directly at S^507-508^ residues, we expressed a recombinant peptide (aa 486 - 536) of KKT2 including or not a mutation of these residues. We demonstrated that recombinant CLK1 could *in vitro* phosphorylate recombinant KKT2 at positions S507-508 (Fig. 4d). Next, we raised a phospho-specific antibody against KKT2^S508^ to follow KKT2 phosphorylation through the cell cycle and after treatment with AB1. The antibody specifically recognises phosphorylation of KKT2^S508^, as phosphorylated KKT2^S508^ was depleted following KKT2 or CLK1 RNAi (Fig. 4e, upper panel), or after treatment with AB1 (Fig. 4e, lower panel; and both endogenous KKT2 and Ty-mNG KKT2 are detected). Phosphorylated KKT2^S508^ was enriched in S-phase after hydroxyurea synchronisation and diminished at 3 hr during anaphase (Fig. 4f). Additionally, we have also compared KKT2 S^508^ phosphorylation in cell cycle sorted parasite populations, observing an increase in the 4C population. Together, these data show that KKT2 phosphorylation is downstream of CLK1 in a kinetochore-specific signalling cascade, and occurs during early metaphase. Considering the strong inhibition correlation (R^2^ = 0.90) between *T. brucei* and *L. mexicana* CLK1 enzyme by AB compound series (as expected, as the two enzymes have 76% sequence identity, Supplementary Fig. 3b,c) and the conservation of KKT2 S508 residue in *Leishmania* and *T. cruzi* (Supplementary Fig. 9) it is quite likely that this signalling pathway is conserved across the trypanosomatids.

In mammals kinetochore assembly is enhanced by mitotic phosphorylation of the Dsn1 kinetochore protein by aurora kinase B, generating kinetochores capable of binding microtubules and promoting the interaction between outer and inner kinetochore proteins ^29^. In T. *brucei*, aurora kinase B (TbAUK1) plays a crucial role in spindle assembly, chromosome segregation and cytokinesis initiation ^23^. Therefore, we asked if TbCLK1 and TbAUK1 are part of the same signalling pathway. We showed that treatment with AB1 does not affect spindle formation (Fig. 3d), in contrast to inhibition of TbAUK1 ^30^. TbAUK1 is a key component of the trypanosome CPC ^31^. To understand if CPC dynamics are impaired by TbCLK1 inhibition, we followed the localisation of TbCPC1 throughout cell cycle before and after AB1 treatment and following TbAUK1 inhibition by Hesperadin ^32^. After treatment with AB1, CPC1 showed a dispersed nuclear pattern that progressively disappeared after nuclear abscission (Fig. 4g middle). This was different to TbAUK1 inhibition by Hesperadin, which prevented trans-localization of the CPC from the spindle midzone, impairing initiation of cytokinesis (Fig. 4g right). Finally, we confirmed that TbAUK1 is not involved in kinetochore assembly since neither KKT2 localisation nor KKT2^S508^ phosphorylation was affected by TbAUK1 inhibition by Hesperadin (Fig. 4h). Recently, a cohort of divergent spindle-associated proteins have been described that are required for correct chromosome segregation in *T. brucei* ^33^. Therefore, we analysed the subcellular localizations of NuSAP1 and NuSAP2 during the cell cycle after TbCLK1 inhibition. NuSAP2 expression in the central portion of the spindle after metaphase release was compromised by TbCLK1 inhibition, whilst NuSAP1 remained unaffected (Supplementary Fig. 10). NuSAP2 is a divergent ASE1/PRC1/MAP65 homolog, a family of proteins that localizes to kinetochore fibres during mitosis, playing an essential role in promoting the G2/M transition ^34^. Considering that NuSAP2 and KKT2 co-localise during interphase and metaphase ^33^, it is likely that KKT2 regulation by CLK1 influences posterior spindle stability and cytokinesis.

Inhibition of Aurora kinase (TbAUK1) does not affect KKT2 S^508^ phosphorylation. <u>Left:</u> KKT2 S^508^ phosphorylation analysed by WB in parasites treated or not with 5x EC_50_ and 2× EC_50_ Hesperadin for 6 hr and 18 hr respectively. Concurrently, AB1 treatment was used as positive control in the same conditions. EF-1 alpha protein expression was used as the loading control. <u>Right:</u> (a) Localisation of TY-mNG KKT2 after 6 hr treatment with 5x EC_50_ Hesperadin. Cells in metaphase are showed. Cells were counterstained with DAPI to visualize DNA (cyan). We next assessed whether KKT2 phosphorylation is required for recruitment of proteins to the trypanosome kinetochore. KKT1 and KKT9 recruitment were impaired in the KKT2^S507A-S508A^ cell line (Fig. 5a), underlining the importance of KKT2 phosphorylation by CLK1 for kinetochore assembly. Individual expression of phospho-mimetics S^507E^ and S^508E^ impaired KKT1 and KKT9 recruitment, but also affected the timing of events during mitosis, with a notable defect in nuclear abscission (Supplementary Fig. 11a). Interestingly, although KKT2 is an essential multi-domain containing protein ^9,35, 36^ and protein kinase activity is required for function (Fig. 4), the localisation of KKT1 and KKT9 to the kinetochore remained unaffected by the loss of KKT2 protein kinase activity (Supplementary Fig. 11b). These data suggest that KKT2 protein kinase activity is required for a function of the kinetochore that is independent from assembly of the complex. In addition to a divergent protein kinase domain, KKT2 contains a divergent polo box domain (PDB), which is sufficient for kinetochore localisation, and a Cys-rich region, found also in KKT3 and KKT20.^36^ It has been proposed that the unique domains structure of kinetoplastid kinetochore proteins is consistent with kinetoplastid kinetochore having a distinct evolutionary origin^9, 36^ and the finding of a unique CLK1/KKT2-centred regulation for kinetochore assembly supports that hypothesis.

Altogether, we propose a model where TbCLK1 progressively phosphorylates KKT2 during S phase, allowing the timely spatial recruitment of the rest of the kinetochore proteins and posterior attachment to microtubules. Inhibition of TbCLK1 activity with AB1 leads to impaired inner kinetochore assembly and irreversible arrest in M phase, suggesting that this defect cannot be repaired by the parasite checkpoint control, implying a dual function of TbCLK1 at different points during chromosome segregation (Fig. 5b). A combination of growth inhibition screening, target identification and characterization resulted in the chemical validation of CLK1 as a pan-kinetoplastid drug target. In addition, the high level of sequence divergence of trypanosome kinetochore complex proteins from humans ^9, 37^ and the unique cell signalling pathways that controls its assembly make the kinetochore a high value target.

## METHODS

### Cell lines maintenance

Bloodstream form of *Trypanosoma brucei brucei* Lister 427, *T. b. gambiense* STIB930 and *T. b. rhodesiense* STIB900 were cultured in HMI-9 media as described elsewhere^1^. Other kinetoplastid parasites *Leishmania donovani* HU3 and *L. mexicana* (MNYC/BZ/62/M379) were cultured in RPMI 1640 media as described earlier ^1, 38^. The *T. cruzi* Tulahuen parasites constitutively expressing *Escherichia coli* β-galactosidase^39^ were maintained in tissue culture as an infection in NIH 3T3 fibroblast cells^1^. NIH 3T3 and human epithelial (HepG2) cells were obtained from ATCC and grown in RPMI media (Life Technologies).

All other recombinant *T. b. brucei* parasites used in this study were derived from monomorphic *T. b. brucei* 2T1 bloodstream forms ^40^ and were cultured in HMI-11 [HMI-9 (GIBCO) containing 10% v/v foetal bovine serum (GIBCO), Pen/Strep solution (penicillin 20 U ml^−1^, streptomycin 20 mg ml^−1^)] at 37 °C/5% CO_2_ in vented flasks. Selective antibiotics were used as follows: 5 μg ml^−1^ blasticidin or hygromycin and 2.5 μg ml^−1^ phleomycin or G418. RNAi or overexpression was induced *in vitro* with tetracycline (Sigma Aldrich) in 70% ethanol at 1 μg ml^−1^. Endogenous Ty, mNeonGreen or myc-overexpression tagging were performed using the pPOTv6 vector ^41^ and pRPa ^40^, respectively. The generation of inducible TbCLK1 RNAi was generated as previously described ^7^. All primers are listed in Supplementary Table 1.

### Determination of growth inhibition activities

All cell based assays were performed as described before ^1, 42^. Briefly, bloodstream form of *T. b. brucei Lister* 427, *T. b. rhodesiense* and *T. b. gambiense* parasites were incubated with varying concentration of compounds for 48 hr. Cell viability was assessed by measuring ATP levels using CellTiter-Glo reagent (Promega) and 50% growth inhibition (EC_50_) was calculated using sigmoidal dose response curves. For both intracellular *T. cruzi* amastigotes and *L. donovani* HU3 amastigotes growth inhibition was measured as described earlier^1^. *L. mexicana* promastigote growth inhibition was assessed by Alamar Blue, whilst intracellular infection was determined after primary peritoneal mouse macrophages were infected with late-log-phase promastigotes at an infection ratio of 10:1; non-internalized parasites were removed by washing the plates with PBS, and cells were cultured with different drug concentrations for 96 hr. Determination of intracellular parasite numbers were done by fixing the cells in methanol and then stained with DAPI. Cytotoxicity (CC_50_) was also measured against mouse fibroblast NIH 3T3 and HepG2 cell lines by incubating compounds for 4 days^1^.

### Overexpression library and viability assay

Single protein kinase overexpression lines (Supplementary methods table 1) were generated by transfecting *T. brucei brucei* 2T1 cells with a tetracycline inducible overexpression plasmid linearized with *Asc*I restriction enzyme. Each overexpression construct contains the open reading frame (ORF) of a single protein kinase in the plasmid pGL2220 (pRPa-iMYCx)^43^. This plasmid integrates at the tagged rRNA spacer (single locus) of 2T1 *T. brucei*.

Protein kinase overexpression cell lines were adjusted to 2 × 10^5^ cells ml^−1^ and induced for 18 hr by the addition of tetracycline to a final concentration of 1 µg ml^−1^ in 70% ethanol. To establish the EC_50_, the protein kinase overexpression cell lines and parental control 2T1 were treated with two-fold increasing concentrations of compounds (with similar DMSO increasing concentration as control). Cell viability was measured at 48 hr with an POLARstar Omega plate reader spectrophotometer; the determination of cell viability was carried out by the established colorimetric technique using Alamar Blue (0.49 mM resazurin in phosphate-buffered saline (PBS)), in a 96-well plate format spectrophotometric assay which measures the ability of living cells to reduce resazurin. We used pentadimine isethionate (Sigma Aldrich) as a positive control. Fluorescence emission was detected using a CLARIOstar ® reader (BMG LABTECH; excitation filter at 540 nm and emissions filter at 590 nm). Fitting of dose-response curves and IC_50_/EC_50_ determination were normalized as percentage of inhibition based on controls. Hesperadin was obtained from ApexBio Technology.

### Immunofluorescence

Cells treated for 6 hr with compounds or DMSO were centrifuged at 1400 g for 10 min before washing twice with TDB-glucose at room temperature. Suspensions were centrifuged at 1000 g for 5 min and pipetted into 6-well microscope slides and dried at RT. Cells were fixed with 25µl 2% paraformaldehyde diluted in PBS and incubated at room temperature for 5 min. Cells were washed in PBS to remove paraformaldehyde prior to washing twice more with PBS and permeabilized with 0.05% NP40 for 10 min. Cells were washed twice in PBS and dried at RT. Mounting media with DAPI was added to each well with a coverslip. Slides were kept at 4 ^o^C before viewing using a Zeiss LSM 880 with Airyscan on an Axio Observer.Z1 invert confocal microscope.

Ty-NuSAP1 and Ty-NuSAP1 were detected by indirect immunofluorescence by using a mouse Imprint ® Monoclonal Anti-Ty1 antibody (clone BB2). Briefly, cells were harvested by centrifugation at 1400 g for 10 min at room temperature, washed, and resuspended in TDB-glucose. 2×10^5^ cells were dried on slides, fixed in 1% paraformaldehyde (PFA) for 1 hr, washed with PBS, blocked with 50% (v/v) foetal bovine serum for 30 min and then incubated with anti-TY (1:800) diluted in 0.5% blocking reagent for 1 hr. Alexa-Fluor® 488 (anti-mouse) was used as secondary antibody (Invitrogen^TM^). Cells were DAPI stained and visualized using a Zeiss LSM 880 with Airyscan on an Axio Observer.Z1 invert confocal microscope.

### Antibodies and western blot

KKT2 and KKT3 phosphorylation profile were analysed by using a SuperSep Phos-tag™ Precast Gel ^27^ according to the manufacturing protocol. Briefly, Ty-mNG KKT2 and Ty-mNG KKT3 were incubated with 5x AB1 EC_50_ for 18 hr and collected for analysis by WB in an EDTA-free RIPA lysis buffer. In parallel, the expression of both proteins were also analysed after 24 hr TbCLK1 RNAi. After electrophoresis, the gel was washed 5 times with 10 mM EDTA transfer buffer to improve transference. Then, the membrane was transfered to a PVDF membrane using a 0.1% SDS Tris-Glycine transfer buffer at 90 mA overnight at 4 ^o^C. The membrane was blocked for 1 hr with 10% BSA and KKT2 and KKT3 phosphorylation pattern was analysed by using an anti-Ty1 antibody. (Supplementary Methods for details)

Anti-phospho KKT2 S^508^ was raised against a synthetic phosphopeptide antigen C-GTRVGS(pS*)LRPQRE-amide, where pS* represent phosphoserine. The peptide was conjugated to keyhole limpet hemocyanin (KLH) and used to immunize rabbits. Phosphopeptide-reactive rabbit antiserum was first purified by protein A chromatography. Further purification was carried out using immunodepletion by non-phosphopeptide resin chromatography, after which the resulting eluate was chromatographed on a phosphopeptide resin. Anti-antigen antibodies were detected by indirect ELISA with unconjugated antigens passively coated on plates, probed with anti-IgG-HRP conjugate, and detected with ABTS substrate. Posterior antigen specificity was confirmed by western blot using KKT2 RNAi and endogenous tagged KKT2 cell lines. Custom antibody was produced by Thermo Fisher Scientific.

## Contributions

J.C.M., T.T.D and S.P.S.R planned the studies. Compounds synthesis, compound docking, PK assays, and library screening by H.X.Y.K, D.P, Y.L.C, S.B.L, S.W, F.S and J.J; HAT animal studies by E.M, R.R, M.P.B and M.K; generation of individual protein kinase cell lines by E.B and M.S; Compound target deconvolution, TbCLK1 functional characterisation, immunofluorescences and data analysis were performed by M.S; protein recombinant production, kinase assays and mass spectrometry by E.F, D.P, C.B-L and M.S; kinetochore foci analysis by M.S, A.J.M.W and M.C.L.;CLK1 and KKT2 recoded and mutants plasmids designed and prepared by M.S, D.P and R.M; J.C.M, S.P.S.R, and M.S prepared and wrote the manuscript. All authors reviewed, edited and approved the paper. J.C.M, S.P.S.R, M.P.B and T.T.D obtained funding.

## Data availability

The data that support the findings of this study are available from the corresponding author upon reasonable request.

## Corresponding author

Correspondence to Jeremy Mottram and Srinivasa P.S. Rao.

## Acknowledgments

This work was supported by the Wellcome Trust (069712). JCM is a Wellcome Trust Investigator (200807). We thank our colleagues in The Bioscience Technology Facility of University of York who provided insight and expertise that greatly assisted our microscopy and flow cytometry research. We thank Gerald Lelais, Vanessa Manoharan, Rima Palkar, Vivian Lim, Christian Noble and Wan Kah Fei for their technical support.

## Supplementary Figure legends

**Supplementary Figure 1.**
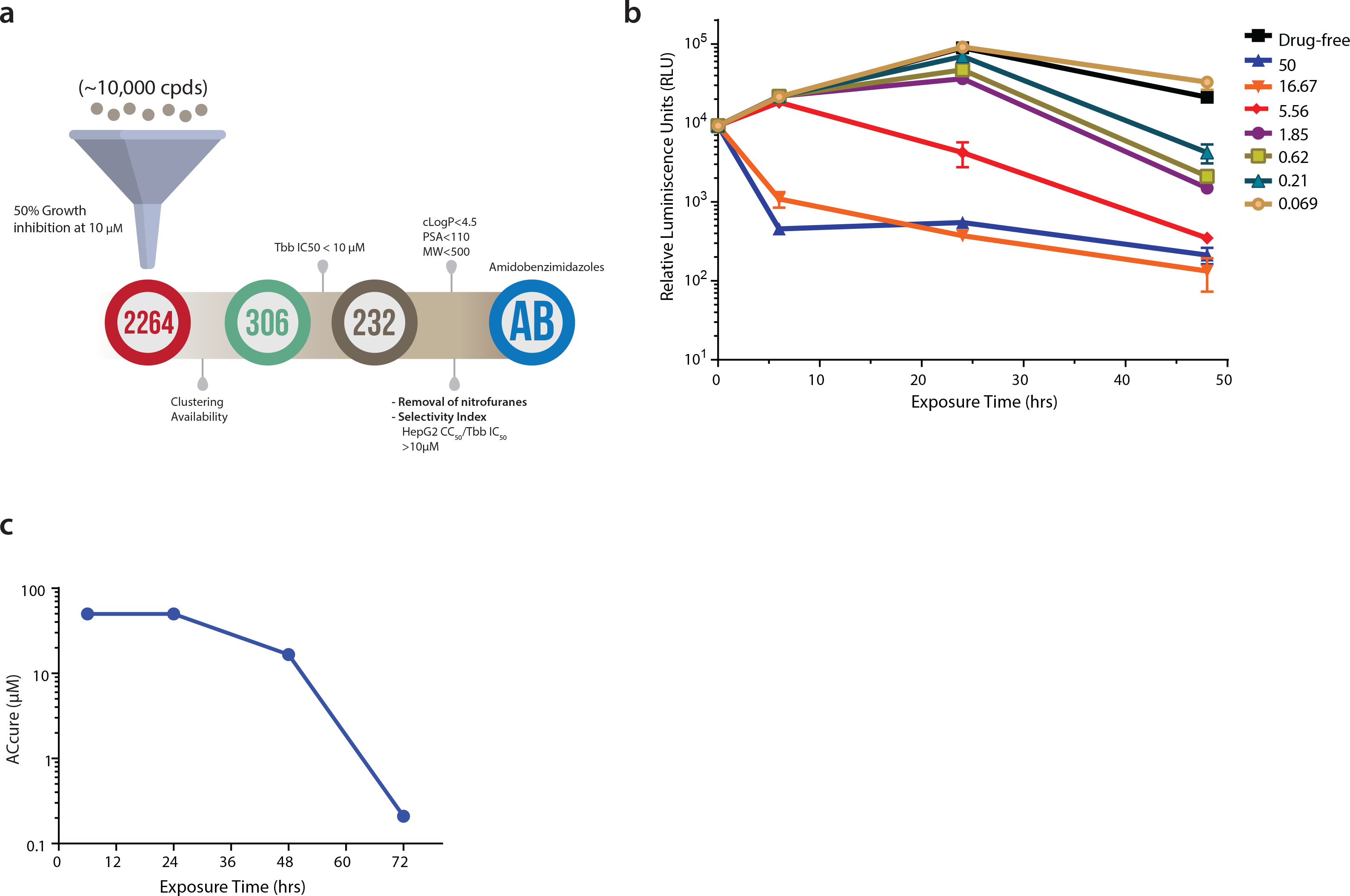
High-throughput screening of kinase library and biological characterization of AB1. (a) Identification of amidobenzimidazole (AB) series as progressable scaffold.
(b) Time to kill analysis of AB1 compound. Note the concentration and time dependent kill of AB1 compound.
(c) The absolute concentrations (AC_cure_) required to achieve sterile cure under *in vitro* conditions with incubation of compound at various time points. Note that as the time of incubation increases, AC_cure_ reduces.

**Supplementary Figure 2.**
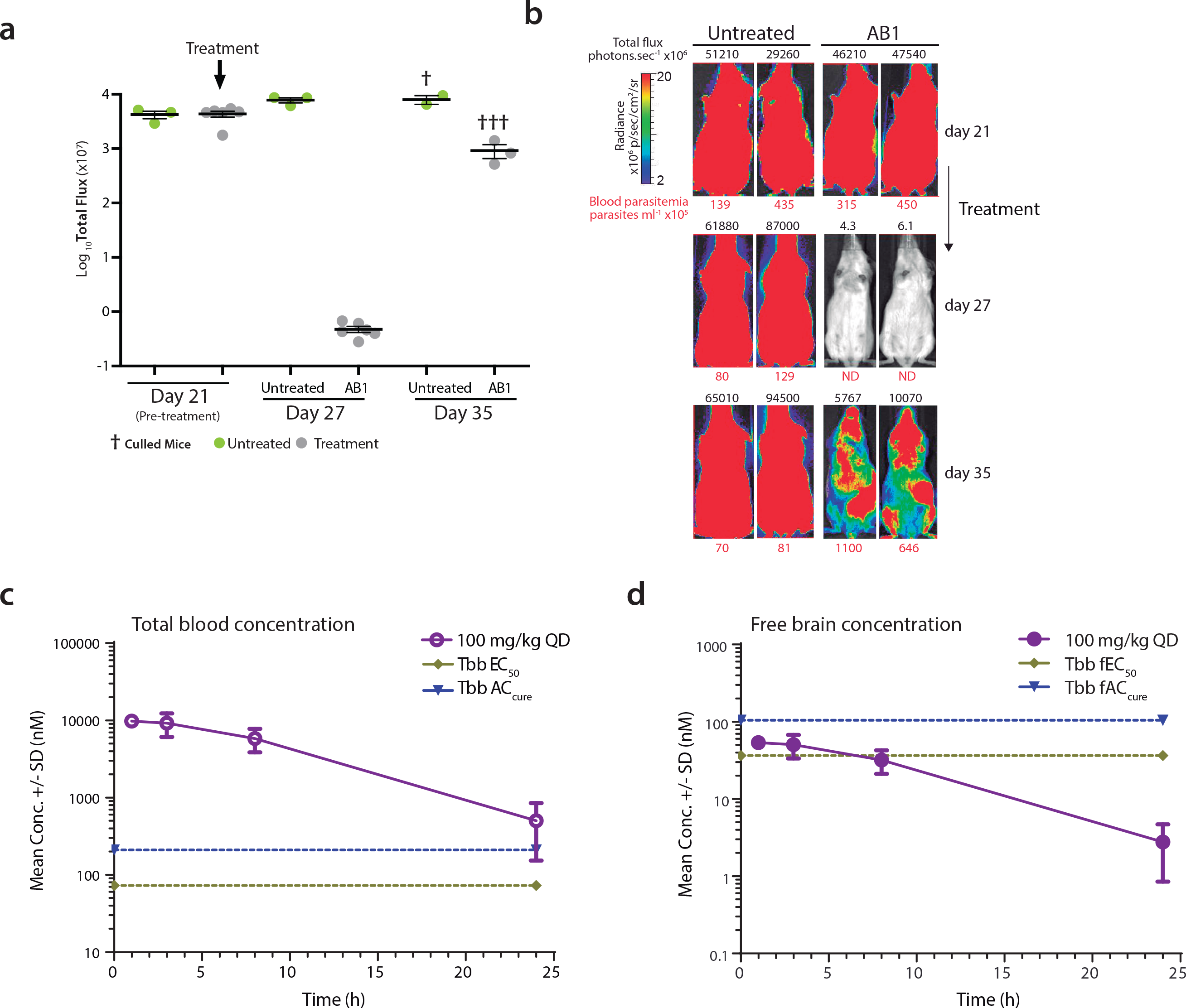
AB1 reduces parasitaemia in a GVR35 strain CNS mouse model of HAT and pharmacokinetics of AB1. (a) Whole body *in vivo* imaging of bioluminescent *T. brucei* before and after AB1 treatment; *Trypanosoma brucei*–infected mice (Day 21) were orally treated with 50 mg kg^-1^ AB1 once-daily for 7 days (n=6 mice, grey) or left untreated (n=3 mice, green); symbols show whole body bioluminescence values for individual mice; mice were euthanized between days 27 and 35.
(b) Images show ventral views of two representative mice of the untreated and AB1-treated groups over the full course of infection. The colour scale indicates bioluminescent radiance in photons/sec/cm^2^/sr. ND, not detected.
(c) Total blood concentration of AB1 in mice at various time points after last dose from the GVR35 strain CNS mouse model HAT. At each time point, 3 mice were bled to collect samples, each point represents mean ± standard error.
(d) Free brain concentrations of AB1 compound in mice calculated by taking into consideration the brain to plasma ratio (0.5), mice plasma protein binding (94%) and rat brain tissue binding (>99%). Note the AB1 concentrations were below *T. b. brucei* EC_50_ and AC_cure_.

**Supplementary Figure 3.**
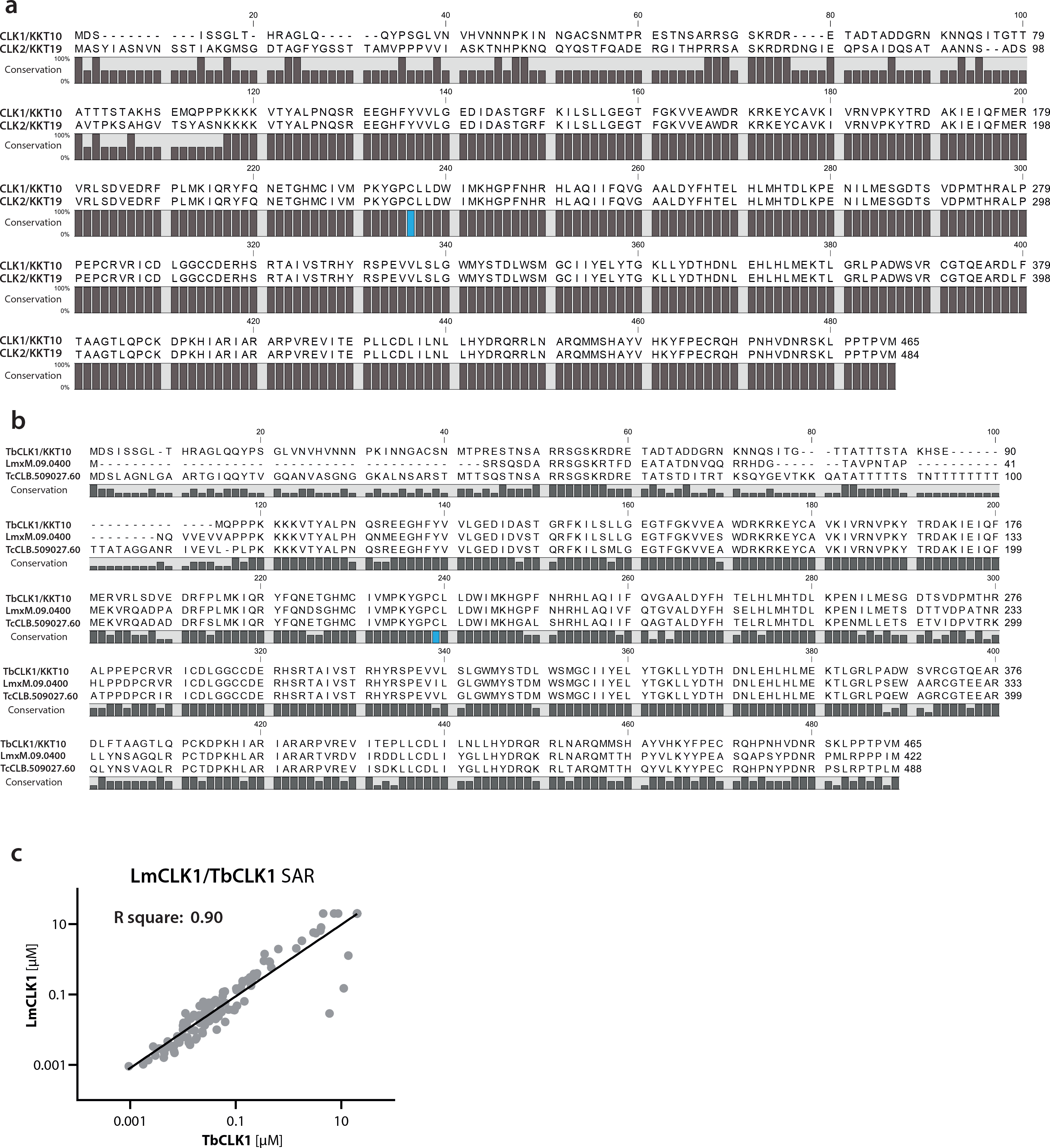
CLK1 and CLK2 protein sequence alignment and CLK1 conservation across trypanosomatids. (a) Alignment of CLK1/KKT10 and CLK2/KKT19 protein sequence in *T. brucei*. The protein sequences of TbCLK1 (Tb927.11.12410) and TbCLK2 (Tb927.11.12420) were aligned in CLC Genomics Workbench 8 and the consensus graph is shown below the corresponding alignment. Residue Cysteine 215 is shown in blue.
(b) The protein sequences of TbCLK1/KKT10 (Tb927.11.12410) and orthologues from *Leishmania mexicana* (LmxM.09.0400) and *Trypanosoma cruzi* (TcCLB.509027.60) were aligned in CLC Genomics Workbench 8. The consensus graph is shown below the corresponding alignment. Residue Cysteine 215 is shown in blue.
(c) Correlation of recombinant kinase inhibition between *T. brucei* and *L. mexicana* CLK1 for the AB series compounds. R square: 0.90

**Supplementary Figure 4.**
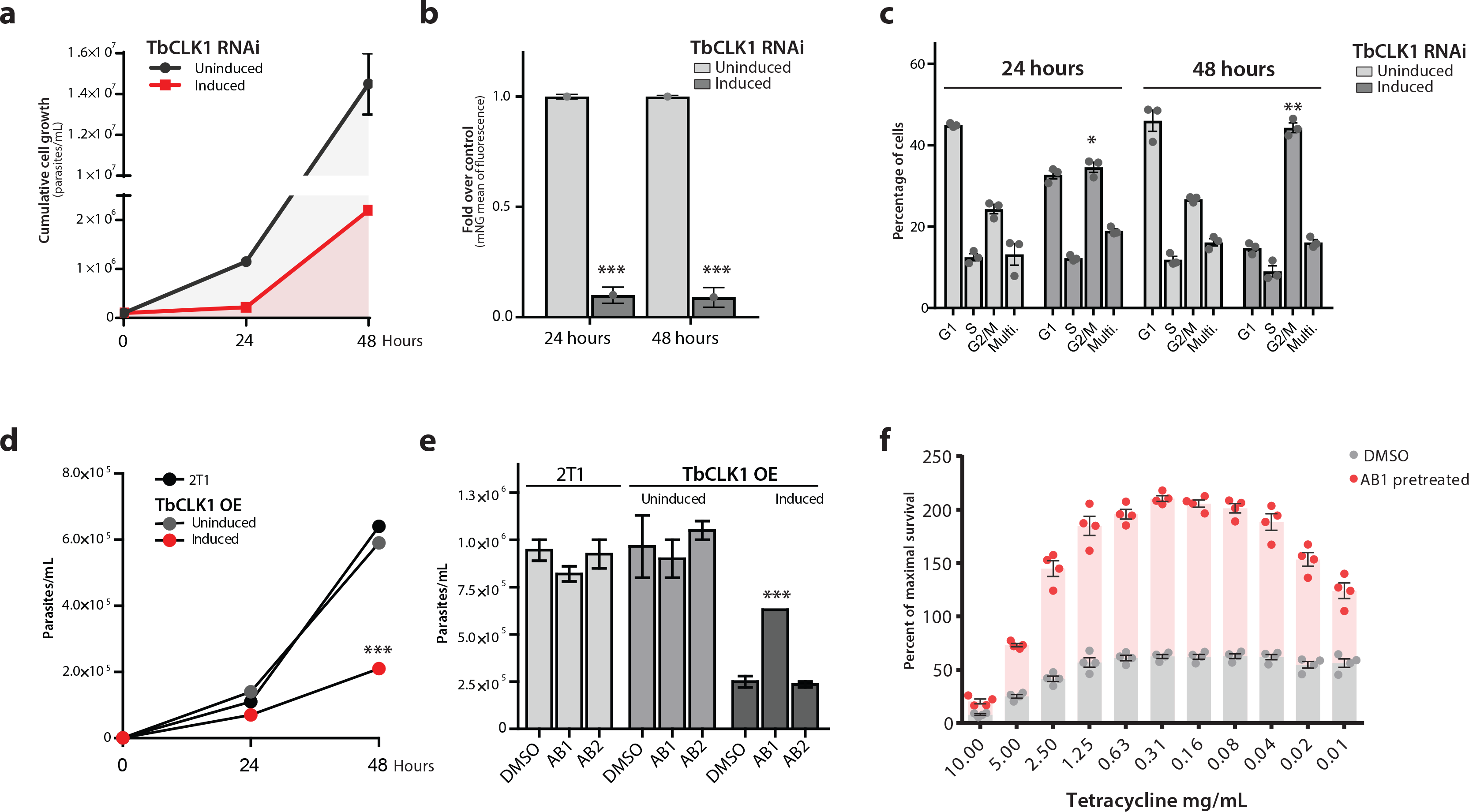
TbCLK1 target validation and TbCLK1 OE toxicity blockage by AB1. (a) TbCLK1 depletion halts cell proliferation. Growth curves of TbCLK1-depleted bloodstream trypanosomes after RNAi induction with tetracycline (in red) is compared with uninduced (in grey) as control. Cultures were diluted daily to 2 × 10^5^ cells mL^-1^ to maintain cell density within a range that supports exponential growth. mNeonGreen (mNG) TbCLK1 endogenously tagged cell line was used to monitor TbCLK1 abundance after RNAi induction by flow cytometry
(b). Error bars, SEM (n=3).
(c) TbCLK1 depletion induces a G2/M cell cycle arrest. Flow cytometry analysis of DNA content in TbCLK1-depleted cells (dark grey) and in the uninduced control (light grey) 24 and 48 h after RNAi induction with 1μg mL^-1^ tetracycline. Error bars, SEM (n=3); P values were calculated using a Two-tailed Student’s t-tests comparing with uninduced control where * p < 0.05, **p < 0.01.
(d-f) TbCLK1 target validation using non-toxic concentrations of AB1 (d) TbCLK1 overexpression impairs parasite growth, (e-f) 24 hr pre-treatment with AB1 non-toxic concentrations (60 nM) reduces the loss of fitness caused by TbCLK1 overexpression induced by tetracycline. Error bars, SEM (n=4); ***p < 0.001 compared to DMSO control.

**Supplementary Figure 5.**
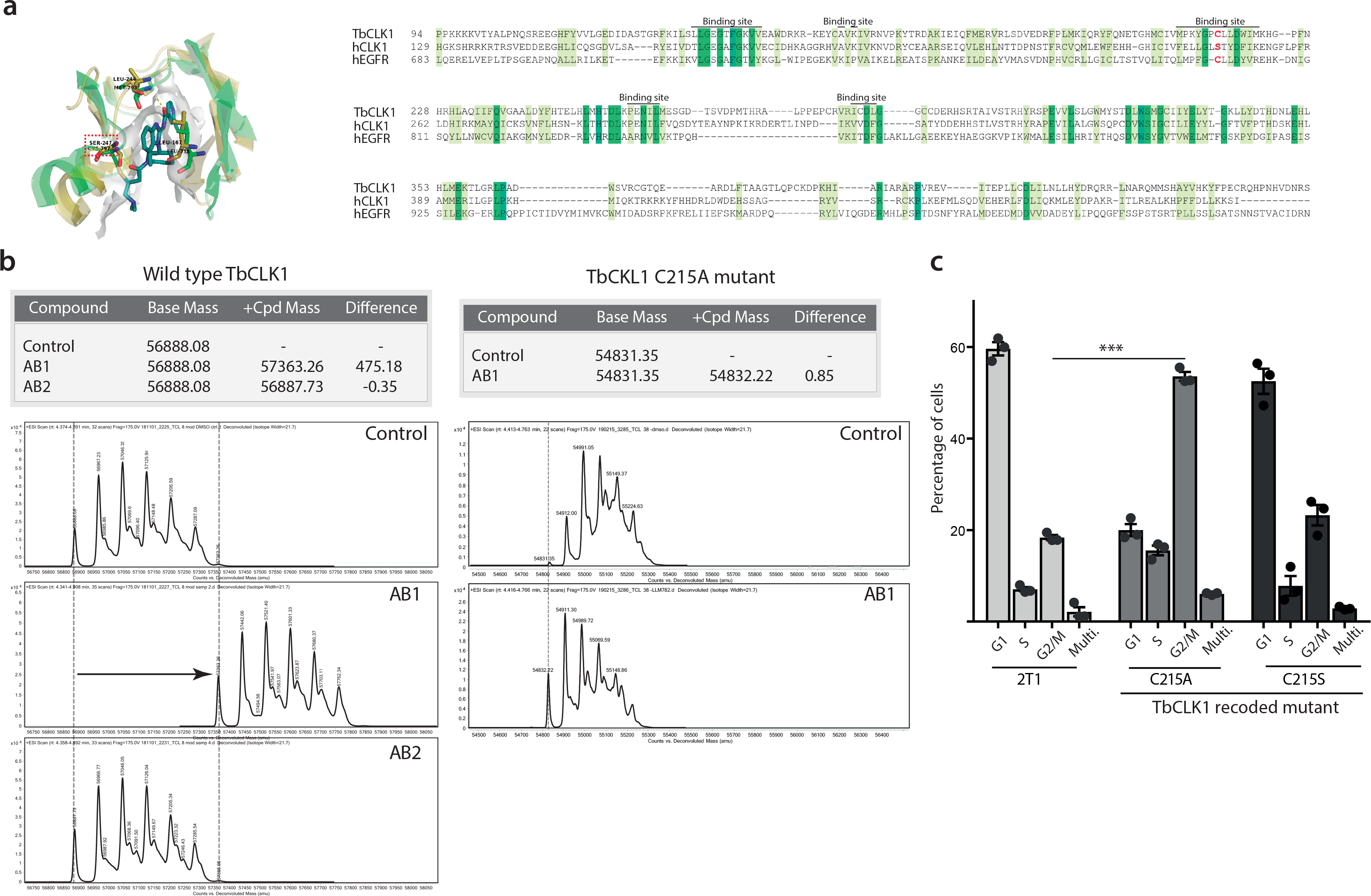
Similarity of hEGFR and CLK1 binding sites highlighting potential of AB1 to covalently bind to TbCLK1. (a) <u>Right:</u> Binding site similarity shown in overlay of hEGFR: AB1 (pdb code: 5feq, green) and hCLK1 with bound ligand deleted (pdb code: 1z57, dark yellow). Key interactions between hEGFR:AB1 and corresponding residues in hCLK1 highlighted in stick. <u>Left:</u> Sequence alignment of hEGFR (Uniprot id: P00533), hCLK1 (Uniprot id: P49759) and TbCLK1 show similarity in binding site residues with presence of covalent cysteine (red) in TbCLK1 explaining selectivity over hCLK1.
(b) Mass spectrum chromatograms of wild type TbCLK1 (left) and TbCLK1 C215A mutant (right) incubated or not with AB1 or AB2 compounds. Y-axis and X-axis represent the counts and Deconvoluted Mass (amu) respectively. Calculated differences after treatment of Bass Mass is showed in the table.
(c) Expression of TbCLK1 C215A results in a G2/M cell cycle arrest. Induction of recoded TbCLK1 C215A and C215S mutants were expressed during 24 hours with tetracycline and cell cycle distribution was determined by flow cytometry and compared with 2T1 parental cell line. Error bars, SEM (n=3).

**Supplementary Figure 6.**
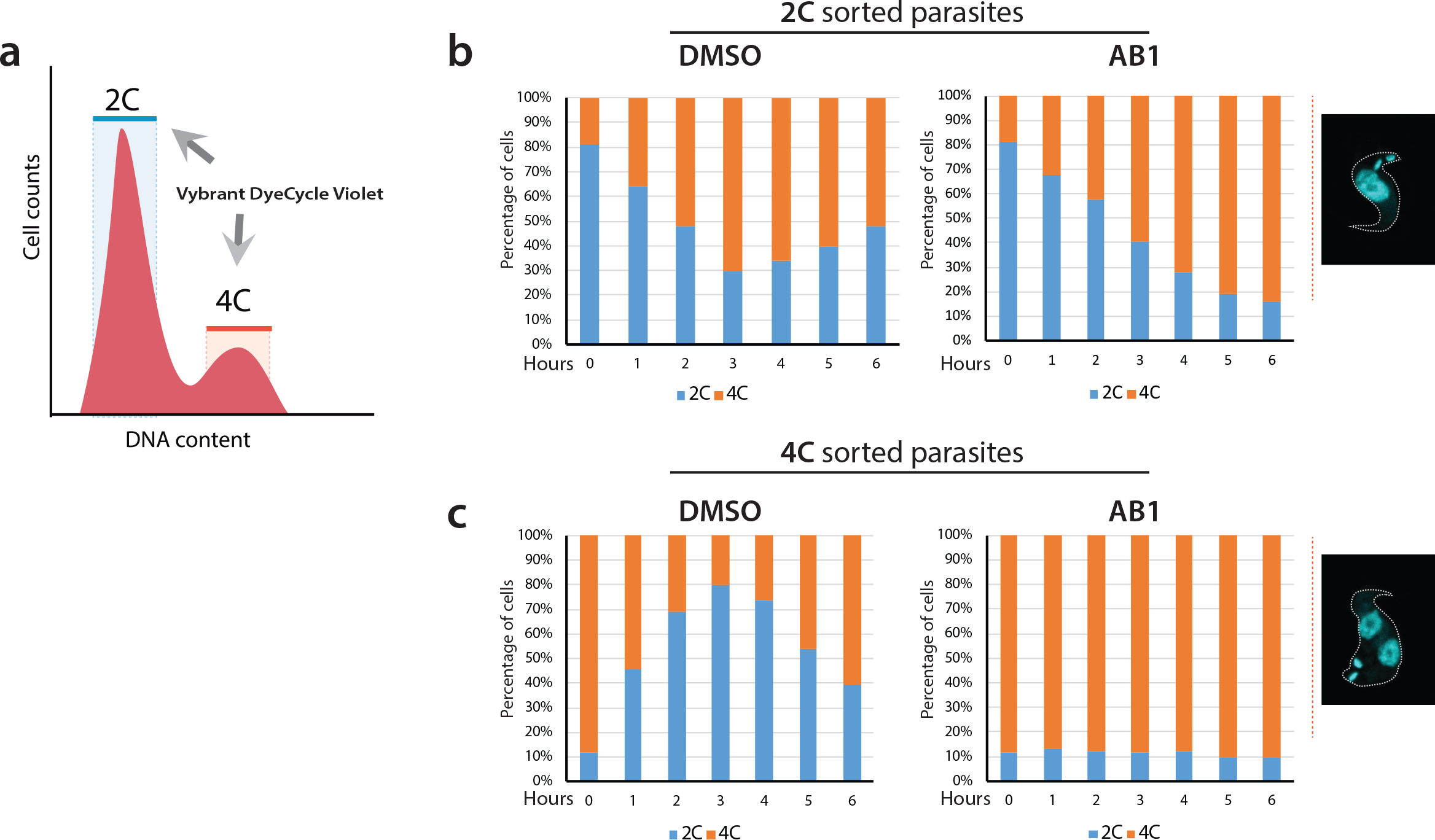
Cell cycle analysis of synchronized *T. brucei* after treatment with AB1. (a) Diagram of 2C (blue) and 4C (orange) cell cycle synchronised bloodstream form *Trypanosoma brucei* using Vybrant DyeCycle Violet-based cell sorting. The selected area shows those cells that were gated for sorting (referred as populations). (b - c) After release, 2C and 4C synchronised parasites were treated or not with 5x EC_50_ AB1. Then, cell samples were collected every hour for DNA flow cytometry analysis of 2C (in blue) and 4C (orange) populations, up to 6 hours.

**Supplementary Figure 7.**
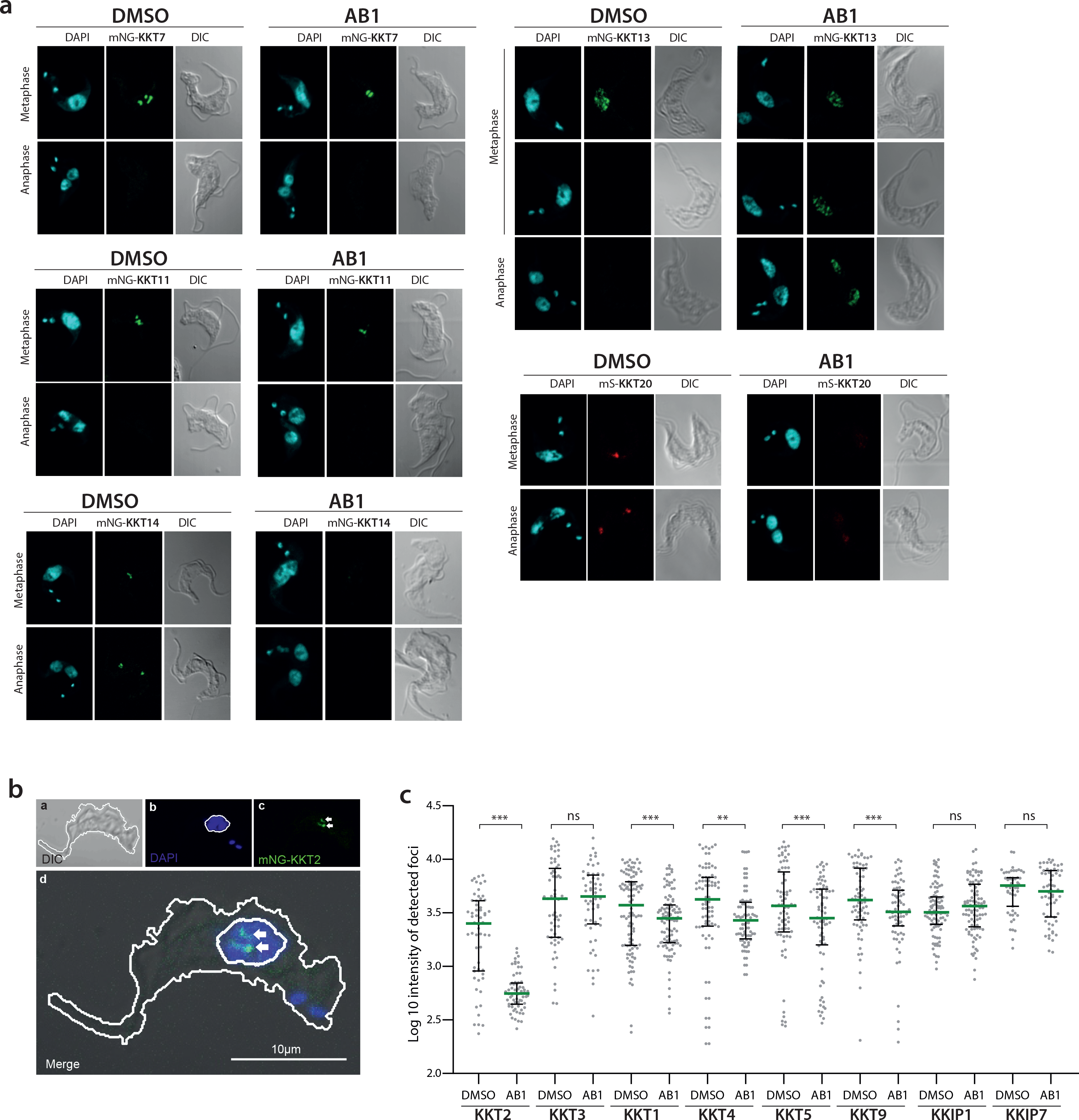
(a) Localization of inner kinetochore core components KKT7, KKT11, KKT13, KKT14 and KKT20 after TbCLK1 inhibition by AB1. Representative fluorescence microscopy micrographs, showing BSF parasites endogenously expressing N-terminal mNeonGreen (mNG) tagged KKTs. For KKT20, BSF parasites endogenously express N-terminal mScarlet (mS) tagged protein. Cells in metaphase and anaphase are shown. Cells were counterstained with DAPI to visualize DNA (cyan).
(b) Graphic representation of strategy used for automated identification of kinetochore and background regions and quantification of fluorescence at kinetochore foci. The region of parasite body and nucleus is masked in white, and the region of interest (ROI) quantified for the kinetochore is highlighted with arrows. In this case, KKT2 foci detection in untreated cells was used as example.
(c) The distribution of kinetochore foci, defined as fluorescence intensities, before and after treatment with AB1. Minima of 60 kinetochore foci were measured for each condition; individual points are shown as grey dots. Median (green) and interquartile ranges (IQR) are shown. ** p<0.01, ***p < 0.001. ns not significant. (Mann–Whitney U test).

**Supplementary Figure 8.**
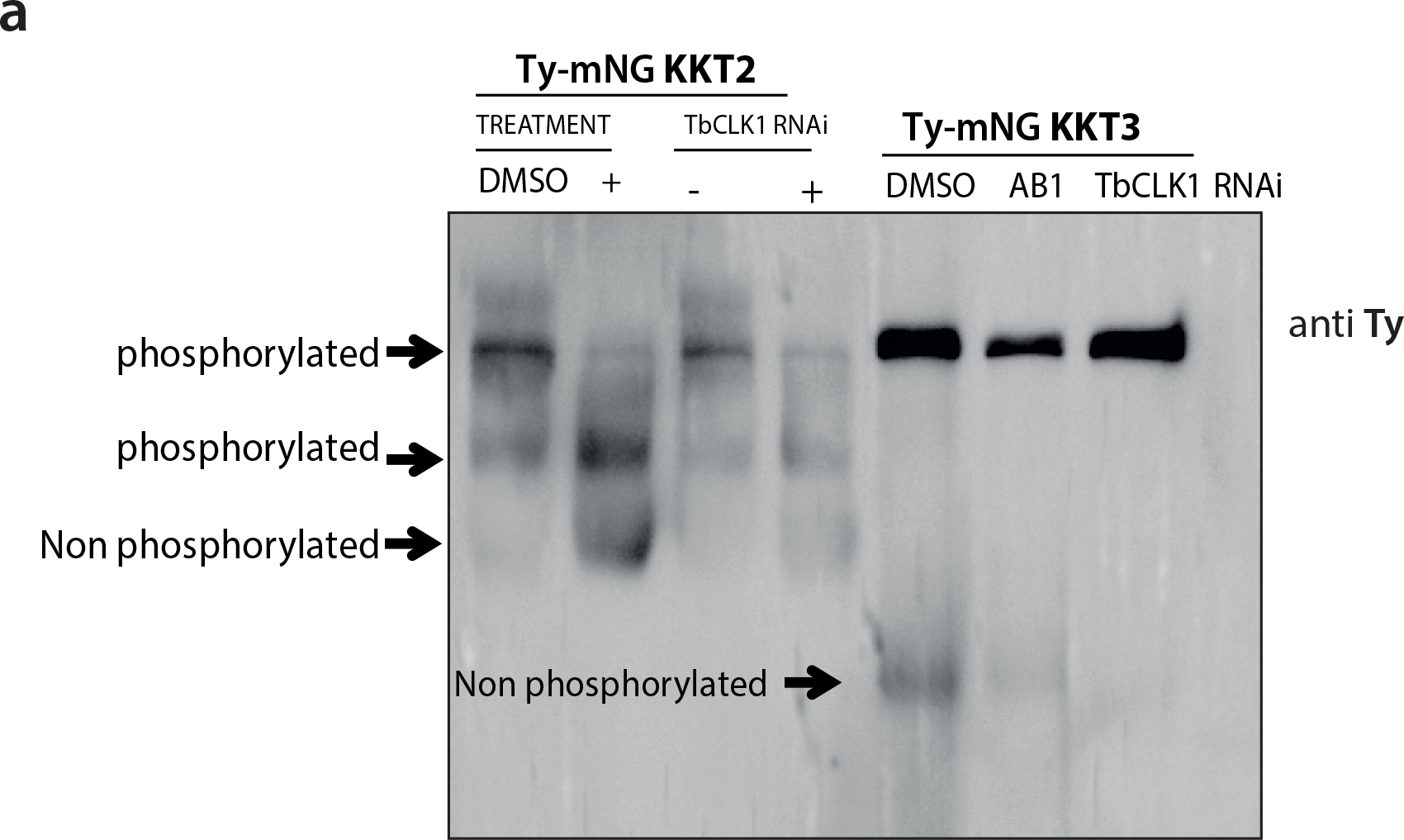
KKT2 and KKT3 phosphorylation pattern after AB1 treatment or TbCLK1 depletion. (a) Phosphorylation pattern of endogenously tagged Ty-mNG KKT2 and Ty-mNG KKT3 cell lines treated or not with 5x EC_50_ AB1 for 6 hr, or after 18 hr of TbCLK1 RNAi induction. Protein samples were collected and resolved by using Phos-tag™ technologies, according to the manufacturer’s protocol^1^.

**Supplementary Figure 9.**
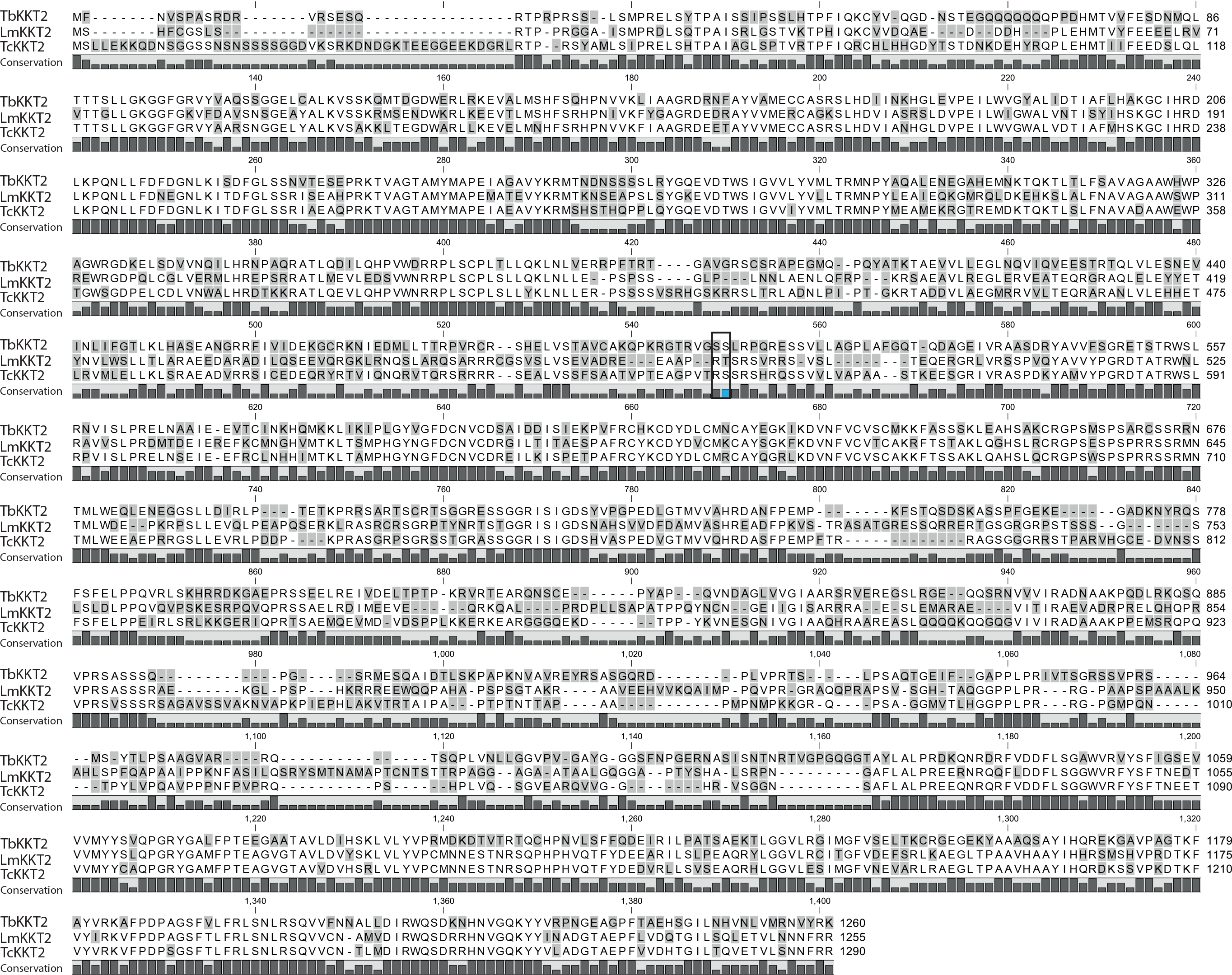
KKT2 S^508^ residue conservation across kinetoplastids. (a) The protein sequences of *Trypanosoma brucei* KKT2 (Tb927.11.10520) and the orthologues from *Leishmania mexicana* (LmxM.36.5350) and *Trypanosoma cruzi* (TcCLB.510285.70) were aligned in CLC Genomics Workbench 8. The consensus graph is shown below the corresponding alignment. KKT2 residues Serine 507 and 508 are shown with a box.

**Supplementary Figure 10.**
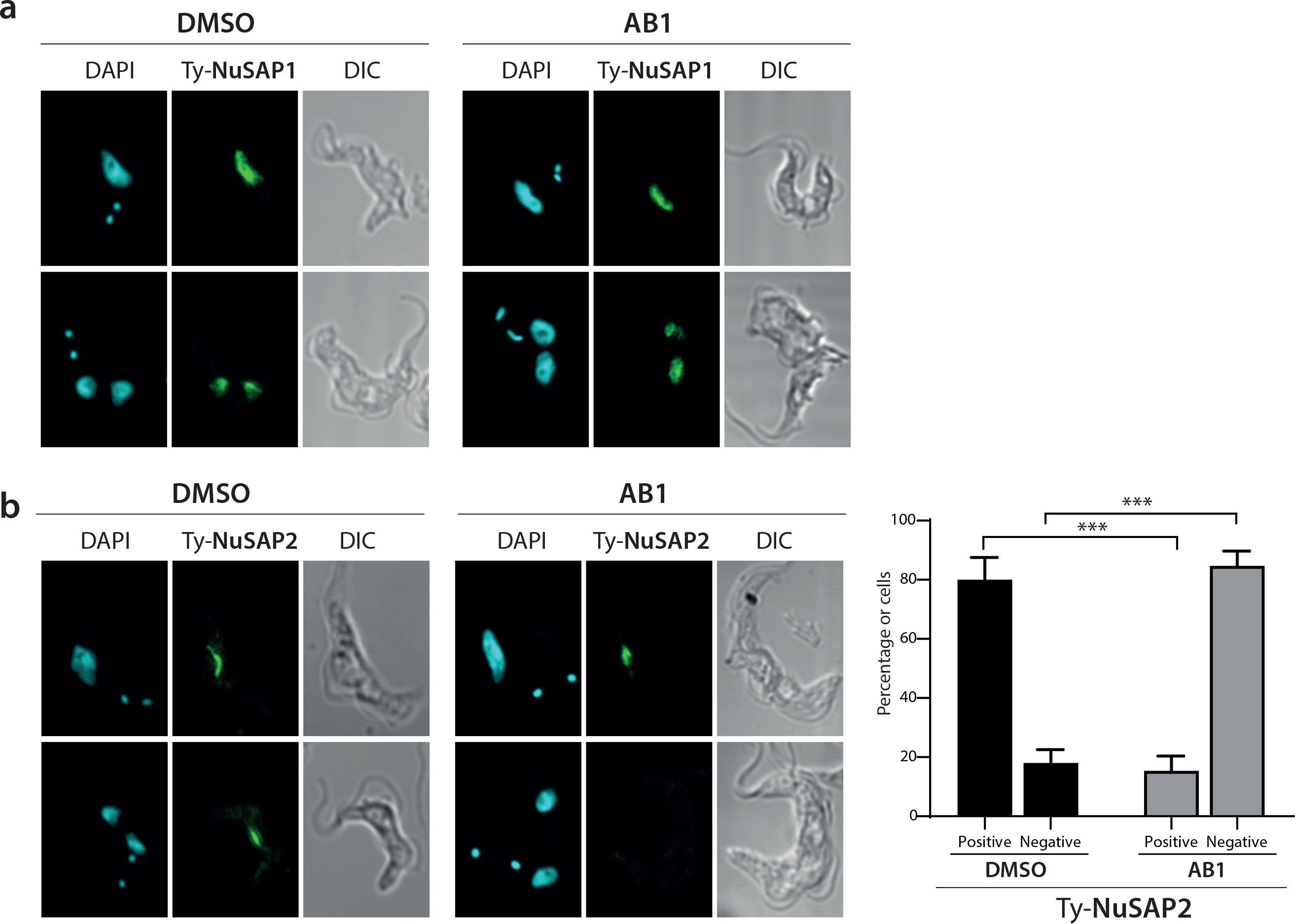
(a) Representative fluorescence microscopy micrographs showing localization of Nucleolar and Spindle Associated Protein 1 (NuSAP1) and 2 (NuSAP2) (b), after TbCLK1 inhibition by AB1. Both proteins were endogenously tagged with a Ty at the N terminus. Cells in metaphase and anaphase are shown. Cells were counterstained with DAPI to visualize DNA (cyan). Lower right panel shows the quantification in percentage of positive or negative expression of NuSAP2 (n=200) during anaphase in control (DMSO) or treated (AB1) parasites. Error bars, SEM; ***p < 0.001 Two-tailed Student’s t-test.

**Supplementary Figure 11.**
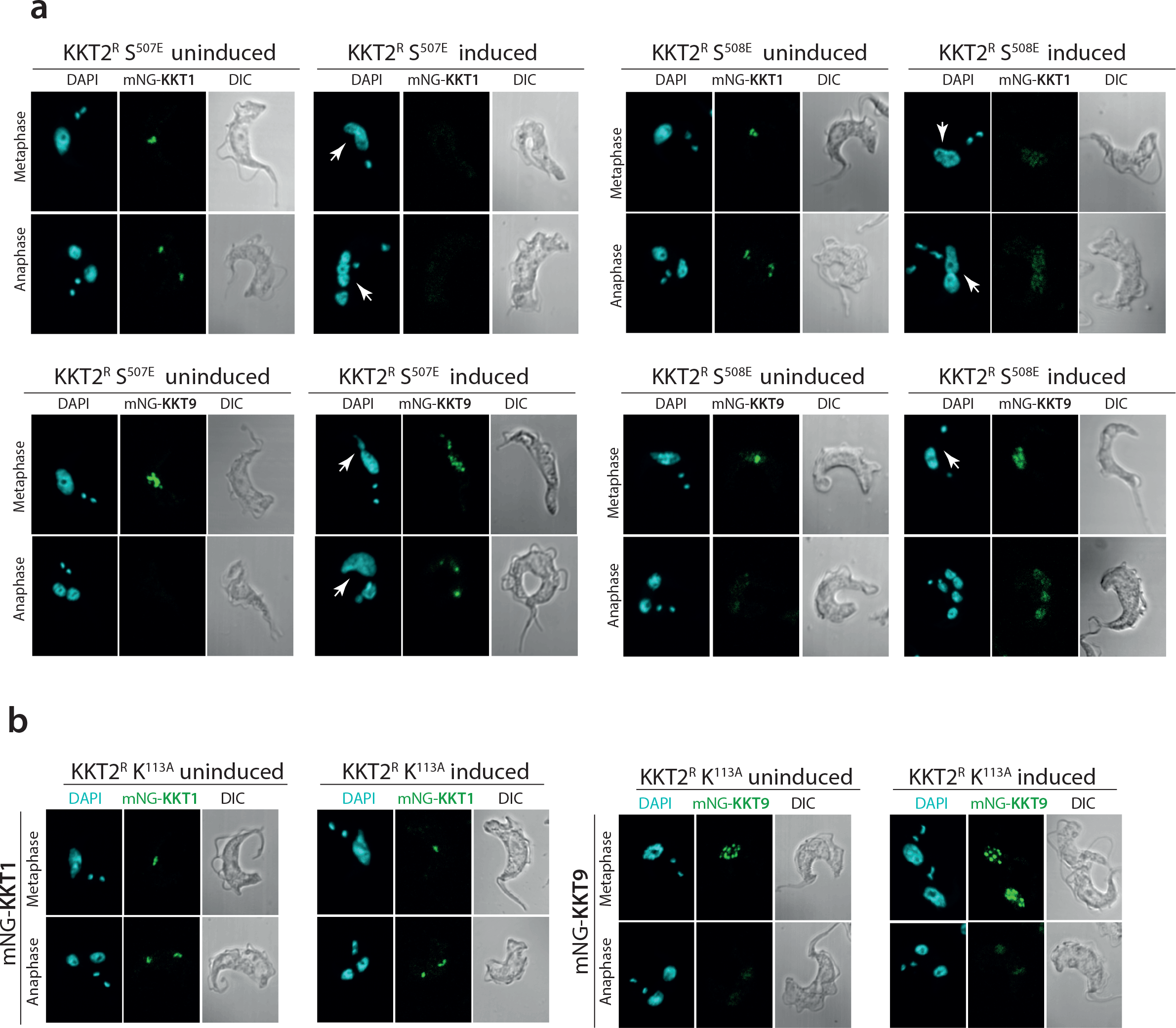
(a) Representative fluorescence microscopy micrographs showing localization of inner kinetochore proteins KKT1 (top panel) and KKT9 (bottom panel) after expression of recoded phosphomimetic KKT2^R^ S^507E^ (left panel) and KKT2^R^ S^508E^ (right panel). In both mutants, KKT1 and KKT9 proteins were endogenously tagged with mNeonGreen (mNG) at the N terminus of KKTs. Abnormal nuclear shape after induction is shown with white arrows. Cells in metaphase and anaphase are shown. Cells were counterstained with DAPI to visualize DNA (cyan). The right panel shows the Nomarsky (DIC) corresponding images.
(b) Localization and expression pattern of inner kinetochore proteins KKT1 (left) and KKT9 (right) after expression of recoded catalytically inactive KKT2^R^ K^113A^. In both mutants, KKT1 and KKT9 proteins were endogenously tagged with mNeonGreen (mNG) at the N terminus of KKTs. Cells in metaphase and anaphase are shown. Cells were counterstained with DAPI to visualize DNA (cyan). The right panel shows the Nomarsky (DIC) corresponding images.

## Supplementary materials and methods

### Protein kinase inhibition assay

Biochemical assays for human EGFR (Epidermal growth factor receptor) and BTK (Bruton’s tyrosine kinase) were carried out using a homogenous time-resolved fluorescence (HTRF) assay as described previously ^1^.

### High throughput phenotypic screening for identification of growth inhibitors.

A kinase focused inhibitor library (containing approximately 10,000 compounds) was screened at a single point concentration of 10 µM for ability to inhibit growth of *T. b. brucei* Lister 427. This resulted in 2264 compounds showing > 50% growth inhibition. Chemi-informatic analysis of hits resulted in ∼200 clusters, representatives of these were subjected to 10 point dose-response growth inhibition assays against *T.b. brucei* Lister 427. Further removal of non-favourable chemical structures such as nitrofuranes, cytotoxicity profiling using the HepG2 cell line (selectivity index of 10) and favourable physico-chemical properties (polar surface area <100; molecular weight <500 and lipophilicity <4.5) led to identification of an amidobenzimidazole scaffold (AB0) for further follow up.

### Determination of solubility, plasma protein binding, brain tissue binding and microsomal clearance.

Solubility of AB0 and AB1 compounds were determined in a high-throughput thermodynamic solubility assay as described previously ^2^. Plasma protein binding was determined for AB1 using mouse blood ^2^, whilst brain tissue binding was determined using rat brain tissues. Intrinsic metabolic clearance of AB0 and AB1 were determined in mouse, rat and human liver microsomes using the compound depletion approach and LC-MS/MS quantification ^3^.

### Archive based structure activity relationship

The Novartis library had several compounds belonging to amidobenzimidazole series in the archive, which were synthesized as a part of other protein kinase inhibitor drug discovery programs. They were further used for structure activity relationship using multiple cellular and enzyme assays as described in this manuscript.

### Kill kinetics and reversibility assay

Ability of AB1 compound to kill *T. b. brucei* Lister 427 was measured over a period of 6, 24 and 48 hr post-compound treatment. The ATP content of parasites was used as a surrogate of the viability of parasites. The assay was conducted similar to the growth inhibition assay stated above with minor modifications. Compound-containing plates were incubated with parasites at 1 × 10^5^ ml^−1^ and at each time point, CellTiter Glo reagent was added to lyse the parasites and luminescence was measured using a Tecan M1000 plate reader after 30 min incubation.

Reversibility assessment to establish time and concentration required to achieve irreversible (relapse-free) growth inhibition under *in vitro* condition was carried out as described elsewhere^4^. The AC_cure_ is the absolute concentration required to achieve sterile cure under *in vitro* conditions with incubation of compound for 72 hr.

### Mouse model of stage I (acute) HAT

The STIB795 acute mouse model mimics the first stage of the disease. Six female NMRI mice were used per experimental group, divided into two cages (A and B). Each mouse was inoculated i.p. with 10^4^ bloodstream forms of STIB795, respectively. Heparinized blood from a donor mouse with approximately 5 × 10^6^ ml^−1^ parasitaemia was suspended in phosphate saline glucose (PSG) to obtain a trypanosome suspension of 1 × 10^5^ ml^−1^. Each mouse was injected with 0.25 ml. Compounds were formulated in 0.5% Tween80 in 0.5% methylcellulose. Compound treatment was initiated 3 days post-infection and administered orally on four consecutive days in a volume of 0.1 ml/10 g. Three mice served as infected-untreated controls. They were not injected with the vehicle alone since we have established in our labs that these vehicles do not affect parasitaemia nor the mice. Blood samples were taken after the 4th treatment. From the mice in cage A, −1 h, 2 h and 8 h after the 4th treatment, 20 uL of blood each were taken from the tail vein and from mice in cage B 1 h, 4 h, and 24 h after the 4th treatment. Parasitaemia was monitored microscopically by tail blood examination twice a week until 31 days post-infection. Mice were considered cured when there was no parasitaemia relapse detected in the tail blood over the 30-day observation period. *In vivo* efficacy studies in mice were conducted at the Swiss Tropical and Public Health Institute (Basel) (License number 2813) according to the rules and regulations for the protection of animal rights (“Tierschutzverordnung”) of the Swiss “Bundesamt für Veterinärwesen”. They were approved by the veterinary office of Canton Basel-Stadt, Switzerland.

### Mouse model of stage II (CNS) HAT

The GVR35 mouse model mimics the second (CNS) stage of the disease. Female CD1 (Charles River UK; ∼8 weeks old; protocol number PPL 60/4442) mice were infected by injection into the peritoneum with 3 × 10^4^ *T. brucei* (GVR35-VSL2) bloodstream form parasites ^5^. Starting on day 21, mice were dosed by oral gavage once-daily with AB1 (n= 6) at 100 mg/kg for 7 days. A group of untreated mice (n= 3) was included as control.

Mice were monitored weekly for parasitaemia from day 21 post-infection. *T. brucei* was quantified in blood samples from the tail vein by microscopy, and *in vivo* bioluminescence imaging of infected mice was performed before treatment on day 21 post-infection and in weeks following the treatment (day 27, 28, 35 post-infection). Imaging on groups of three mice was performed 10 min after i.p.injection of 150 mg D-luciferin (Promega)/kg body weight (in PBS) using an IVIS Spectrum (PerkinElmer) as described previously ^6^. Mice were euthanized between days 27 and 35 by cervical dislocation. Data analysis for bioluminescence imaging was performed using Living Image Software. The same rectangular region of interest (ROI) covering the mouse body was used for each whole body image to show the bioluminescence in total flux (photons per second) within that region. Image panels of whole mouse bodies are composites of the original images with areas outside the ROI cropped out to save space.

All animal procedures were undertaken in adherence to experimental guidelines and procedures approved by The Home Office of the UK government. All work was covered by Home Office Project Licence PPL60/4442 entitled “Molecular Genetics of Trypanosomes and Leishmania”. All animal protocols received approval from the University of York and University of Glasgow Ethics Committees.

### Pharmacokinetic analysis of AB1

Blood samples from mice were collected at 1, 3, 8 and 24 hour post last dose of CNS stage mice studies. Total concentration of AB1 compound in blood was analysed using standard LC-MS/MS as described elsewhere ^1^. Free concentration in brain was calculated by taking in to consideration the brain to plasma ratio, plasma protein binding and brain tissue binding.

### Cell cycle analysis and Cell Sorting

Bloodstream form *T. brucei* cell lines were incubated or not for 6 hr with AB compounds at a final concentration of 5X the individual EC_50_ value for each compound (averaged from viability assays). Control cultures were treated with 0.5µl DMSO. Cultures were pelleted and cells were collected and washed once in *Trypanosoma* dilution buffer (TDB) supplemented with 5 mM of EDTA and resuspended in 70% methanol. Cells were centrifuged at 1400 g for 10 min to remove methanol and washed once in TDB 1x with 5mM EDTA. Cells were resuspended in 1ml TDB 1x with 5mM EDTA, 10µg ml^−1^ of propidium iodide and 10µl of RNase A. Cell suspensions in 1.5 ml tubes were wrapped in foil to avoid bleaching by light. Cells were incubated for 30 min at 37°C in the dark until FACS analysis. Cells were analysed for FACS using a Beckman Coulter CyAn ADP flow cytometer (excitation; 535, emission; 617).

In the cell cycle analysis, TbCLK1 OE was induced during 18 hr with tetracycline (1 μg ml^−1^) and later treated with 4X the individual EC_50_ value for each compound for 4 hr (maintaining tetracycline induction), and finally collected for flow cytometry as above.

Parasite cell sorting was conducted as described previously ^7^. Briefly, cell lines were harvested during exponential growth by centrifugation for 10 min in a clinical centrifuge at room temperature. The parasite pellet was then resuspended at a concentration of 1 × 10^6^ cells ml^−1^ in HMI-9 medium supplemented with 2% FCS and 10 μg ml^−1^ penicillin/streptomycin. Vybrant DyeCycle Violet (Molecular Probes, Invitrogen) was added to a final concentration of 1 μg ml^−1^and the cell suspension incubated for 30 min at 37 °C, the tube being protected from light by wrapping in aluminium foil. The samples were then centrifuged and resuspended back in the staining media prior to sorting on a MoFlo XDP Sorter (Beckman Coulter Life Sciences). During and after the sorting, the samples were cooled to below 20 °C to limit cell metabolic activity. The dye was excited using a 407 nm Violet laser and emission detected via a 450/40 bandpass filter. Live parasites were gated based on FSC/SSC profiles, and the gates were set up to collect only the 2C fraction (G0/G1 cells) and 4C fraction (G2, mitotic and post-mitotic cells), to ensure efficient discrimination and selection of these cell-cycle stages.

Hydroxyurea-induced synchronisation of cell lines was obtained by incubating parasites in exponential growth phase with 10 µM of Hydroxyurea (HU) (Sigma Aldrich) for 6 hr. Removal of HU from the culture medium was achieved by centrifuging cells at 1400 g for 10 min, washing twice with fresh (drug free) medium and resuspending cells in medium lacking HU. Subsequently, samples were collected each hr for posterior cell cycle analysis by propidium iodide staining.

### Recombinant assays and enzyme purification

Full-length TbCLK1 (Q382U0) and LmCLK1 (E9AMP2) CDS were cloned in pET28a PreSc-His and pET24-MBP-TEV vector respectively. Human CLK1 (hCLK1; P49759) was obtained from Promega. Recombinant expression was carried out by lactose autoinduction in Terrific Broth containing 0.4% glycerol, 0.05% glucose, 0.05% lactose, 0.05% arabinose and buffered by 100 mM sodium phosphate (pH 7.0). In brief, 0.7 L of this media (in a 2.8 L Fernbach flask) was inoculated at 0.1 OD600 with an overnight Luria Broth culture and shaken at 37 °C and 250 rpm for 2.5 hr. Then, temperature was lowered to 18 °C and the culture was allowed to grow and induced overnight and harvested 20-24 hr later. Cells are pelleted and stored at −80 °C prior to purification. Cell lysis was done by sonication in an ice bath (20 sec ON/OFF, 3 min active sonication at 70-110 watts power) in 40 mL equilibration buffer (25 mM HEPES pH 7.5 300 mM NaCl 5% glycerol 0.5 mM TCEP) and the clarified lysate is purified by IMAC on a 5 mL HisTrap column (GE Healthcare). The IMAC elution was further purified by sizing on a 300 mL Superdex 200 prep grade column (GE Healthcare) packed in a 2.6 cm diameter housing. Included volume fractions were pooled and analysed by SDS-PAGE or LC-MS.

### Structure–activity relationship (SAR) of AB series compounds

rTbCLK1, rLmCLK1 and hCLK1 enzyme activity assays were performed in white 384 well, solid bottom, Small Volume™ plates (GREINER). The assay buffer contained 40 mM Tris (pH 7.5), 20 mM MgCl_2_, 0.1mg/ml BSA and 2 mM DTT. Each enzyme (TbCLK1 and LmCLK1= 3 nM; hCLK1 = 50nM) was first incubated with different compound serial dilutions or DMSO control during 10 min and then ATP (10 µM) and MBP substrate (0.1 mg ml^−1^) (Myelin basic protein, Dephosphorylated, Sigma-Aldrich Catalogue no: 13-110, LOT: 3107375) mixture was added to initiate the reaction. After 35 min reaction at room temperature, the ADP-Glo reagent and detection solution were added following the technical manual of ADP-Glo^™^ kinase assay kit (Promega). The luminescence was measured on CLARIOstar BMG LABTECH microplate reader. In all the assays, staurosporine was used as positive control of inhibition. IC_50_ values were plotted using GraphPad Prism software.

### Multiple sequence alignment

The Uniprot knowledgebase ^8^ and TriTryDB ^9^ were used to retrieve fasta sequence files for the sequences of hEGFR (P00533), hCLK1 (P49759) and TbCLK1 (Q382U0). The multiple sequence alignment was performed using the Needleman-Wüncsh algorithm ^10^ modified to include zero end gap penalties (ZEGA) ^11^ with the Gonnet substitution matrix ^12^, as implemented in ICM molecular modelling package (ICM version 3.8-6a, Molsoft LLC).

### Structural alignment of hEGFR and hCLK1

To compare the three-dimensional similarity and assess the potential for the AB1 ligand to bind to CLK1, the three-dimensional coordinates from the hEGFR:AB1 cocrystal (PDB code 5feq) was aligned to hCLK1 (PDB code: 1z57). The ICM modelling suite was used to calculate the structural alignment for the pair using the Cα coordinates of the backbone atoms, resulting in high similarity between the 3D fold of the proteins in the region of the binding site.

The sequence and structural similarity contributed towards evidence of AB1 potentially binding to hCLK1 and aided the selection of mutations which would affect the binding and thus confirm the binding of AB1 in TbCLK1.

### Analysis of covalent modification by LC-MS

2 µM of WT TbCLK1 and TbCLK1 C215A were incubated with 4 µM compound at RT for 1 hr, and quenched with 80 µM DTT. Approximately 18 pmol of protein is loaded onto a 2.1 × 50 mm 1000Å PLRP-S column (Agilent) equilibrated in 5% buffer B and eluted by a 2 minute 5-65% gradient at 0.2 mL/min into an Agilent 6530 QToF equipped with an electrospray source. <u>Buffer A</u> is 0.1% formic acid in water, and <u>Buffer B</u> is 0.1% formic acid in acetonitrile. Source voltage was 5500V, drying gas was 12 L/min at 350 C, and the nebulizer was at 60 psig. Fragmentor, skimmer and octopole are at 175 V, 60 V and 750 V, respectively. Mass spectra are acquired at 1 Hz over the m/z range 400-2000. An averaged m/z spectrum representing the total ion chromatogram peak of the protein is extracted, and a zero-charge mass spectrum is generated by maximum entropy deconvolution is performed over a range of 6,000-225,000 Daltons with a 0.5 Da step.

### Kinetochore foci intensity measurement

Three channel image stacks of mNeonGreen labelled kinetochore components in fixed trypanosomes were analysed using bespoke Matlab software (available here https://github.com/awollman). Blue, nuclear stained images were first segmented by thresholding using Otsu’s method and applying a series of morphological transformations to remove holes and any objects smaller than 300 pixel area. This allowed the nucleus to be segmented and removed any detected mitochondria, also stained by DAPI. The whole cell was then segmented from the DIC image, using edge detection and similar morphological transformation, combined with watershedding, using the nuclear mask as ‘seeds’ for each cell. Finally, bright foci were detected in the mNeonGreen image using spot detection software (available here https://awollman.github.io/single-molecule-tools/), optimised for detecting and characterising low intensity foci in noisy cellular environments ^13, 14^. In brief, candidate foci were detected by thresholding and Gaussian masking, before their local background corrected intensity was determined and accepted if above a threshold based on the standard deviation of local pixel noise. Each detected cell was assigned a tracking number and foci categorised into each cell. This allowed for manual assignment into cell cycle stage.

### Recoded Plasmids

Recoded KKT2 was synthesised by Dundee cell products. The recoded KKT2 sequence (KKT2^R^) codes for the same amino acid sequence as KKT2 but only shares 94.23% nucleotide identity. All segments of identity between KKT2 and KKT2^R^ are less than 20 base pairs long. KKT2*^R^* was inserted into the plasmid pGL2243 using *Xba*I and *Bam*HI restriction sites, generating pGL2492. This plasmid is designed to constitutively express KKT2 from the tubulin locus, with the addition of a C-terminal 6x HA tag. To express catalytically inactive KKT2 and phospho-mutants, the active site lysine (K^113^) and serine (S^5^, S^8^, S^507-S508^ and S^828^) were changed to alanine by mutating pGL2492, carrying the coding sequence for KKT2, using site directed mutagenic PCR as follows:

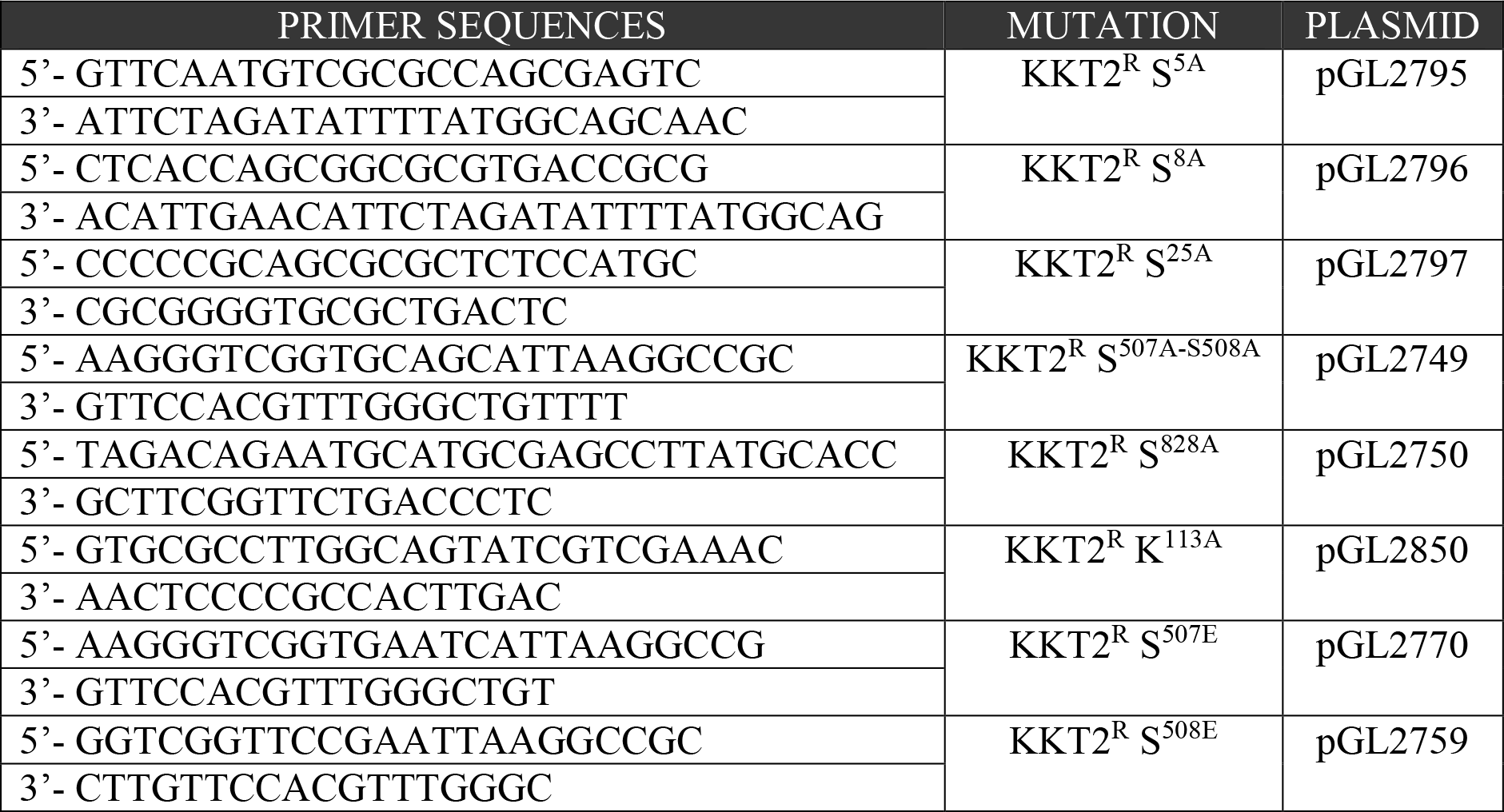

Recoded CLK1 was synthesised by Eurofins Genomics. The recoded CLK1 sequence (CLK1^R^) codes for the same amino acid sequence as CLK1 but only shares 95.06% nucleotide identity. All segments of identity between CLK1 and CLK1*^R^* are less than 20 base pairs long. CLK1^R^ was inserted by Gibson assembly® (New England Biolabs) into the plasmid pGL2492 using *Xba*I and *Bam*HI restriction sites, generating pGL2832. This plasmid is designed to constitutively express TbCLK1 from the tubulin locus, with the addition of a C-terminal 6x HA tag. To express the cysteine 215 mutants, the cysteine 215 was changed to serine by mutating pGL2832, carrying the coding sequence for TbCLK1, using site directed mutagenic PCR as follows:

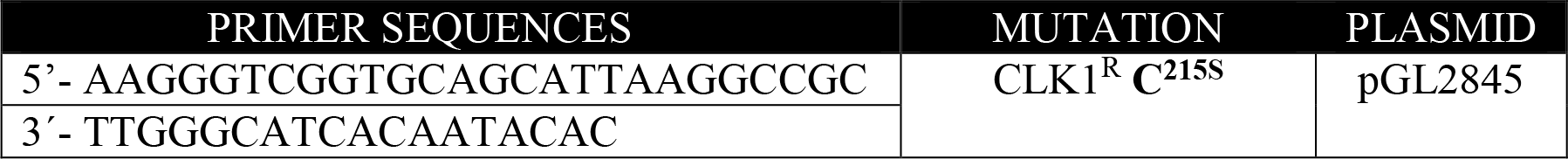

### Antibodies and western blot

Parasites were washed with trypanosome dilution buffer (TDB) supplemented with 20 mM of glucose. After centrifugation, the samples were resuspended in the RIPA buffer (New England Biolabs) supplemented with protease and phosphatase inhibitors obtained from Promega and Roche Life Science respectively. All samples were quantified by *Bradford protein* assay (Bio-Rad), 25 µg of protein was loaded, resolved in a 4-20% NuPAGE Bis-Tris gel (Invitrogen) in NuPAGE MOPS running buffer and transferred onto Hybond-C nitrocellulose membranes (GE Healthcare) at 350 mA for 2 hr or, for high molecular weight proteins, overnight at 4°C.

After transfer, membranes were washed once in 1x TBST (tris-*buffered* saline (TBS), 0.01% Tween-20 (Sigma Aldrich)) for 10 min then incubated for 1 hr in blocking solution (1x TBST, 5% BSA) or, if required, overnight at 4°C. Next, the membrane was rinsed for 10 min in 1X TBST and placed in blocking buffer containing the required primary antisera for 1 hr at room temperature or overnight at 4°C. The membrane was then washed 3 times with TBST and placed in blocking solution containing the appropriate fluorescent secondary antisera for 1 hr.

| Antibody | Obtained from |
| --- | --- |
| Mouse Imprint Monoclonal **Anti-Ty1** antibody (clone BB2) | Sigma Aldrich (SAB4800032) |
| Mouse Anti-HA (clone 12CA5) | Roche (11583816001) |
| Mouse Anti-EF1α Antibody, (clone CBP-KK1) | Merck-Milipore (05-235) |
| Rabbit Anti-phospho KKT2 S<sup>508</sup> | Invitrogen |
| StarBright™ Blue 520 Goat anti-Mouse IgG | BIORAD (12005867) |
| StarBright™ Blue 700 Goat Anti-Mouse/Anti-Rabbit IgG | BIORAD (12004162) |

### General Statistics.

All statistical analysis was performed using GraphPad Prism 8 (http://www.graphpad.com/scientific-software/prism/). The appropriate tests were conducted and are as detailed in the corresponding figure legends.

